# Polyploidy, regular patterning of genome copies, and unusual control of DNA partitioning in the Lyme disease spirochete

**DOI:** 10.1101/2022.07.05.498848

**Authors:** Constantin N. Takacs, Jenny Wachter, Yingjie Xiang, Xheni Karaboja, Zhongqing Ren, Molly Scott, Matthew R. Stoner, Irnov Irnov, Nicholas Jannetty, Patricia A. Rosa, Xindan Wang, Christine Jacobs-Wagner

## Abstract

*Borrelia burgdorferi*, the tick-transmitted spirochete agent of Lyme disease, has a highly segmented genome with a linear chromosome and various linear or circular plasmids. Here, by imaging several chromosomal loci and 16 distinct plasmids, we show that *B. burgdorferi* is polyploid during growth in culture and that the number of genome copies decreases during stationary phase. *B. burgdorferi* is also polyploid inside fed ticks and chromosome copies are regularly spaced along the spirochete’s length in both growing cultures and ticks. This patterning involves the conserved DNA partitioning protein ParA whose localization is controlled by a potentially phage-derived protein, ParZ, instead of its usual partner ParB. ParZ binds its own coding region and acts as a centromere-binding protein. While ParA works with ParZ, ParB controls the localization of the condensin, SMC. Together, the ParA/ParZ and ParB/SMC pairs ensure faithful chromosome inheritance. Our findings underscore the plasticity of cellular functions, even those as fundamental as chromosome segregation.

## INTRODUCTION

Lyme disease is the most prevalent vector-borne infectious disease in North America and Europe^1^. Its geographic range has steadily spread over the years, with caseloads recently estimated to be near 500,000 per year in the United States^1,2^. Lyme disease is caused by *Borrelia burgdorferi* and related spirochete bacteria^3^. In nature, Lyme disease spirochetes undergo a transmission cycle between *Ixodes* hard tick vectors and warm-blooded vertebrate hosts^4^. Infection in humans via a tick bite can result in a wide variety of symptoms when left untreated. Disease manifestations range from skin rashes, fever, and malaise during early stages of the disease to arthritis, carditis, and neurological symptoms during later stages^3^. Given *B. burgdorferi*’s medical relevance, we set out to study basic biological processes necessary for cell proliferation. This topic is of considerable interest because bacterial multiplication is a prerequisite for successful transmission, host infection, and disease causation.

*B. burgdorferi* was identified in 1982^5^. Despite four decades of research, many of the fundamental cellular processes underlying the ability of this bacterium to self-replicate remain understudied. This is in part because *B. burgdorferi* has a long doubling time (5 to 18 h) in culture^6–11^. Additionally, genetic manipulation of this organism, while possible^12–14^, remains challenging compared to that of *Escherichia coli* and other common model bacteria. However, knowledge obtained from the study of model bacteria does not always translate to unrelated species, including spirochetes. These bacteria form a phylum of particular interest because, in addition to *B. burgdorferi*, it includes important human pathogens such as the agents of relapsing fever, syphilis, and leptospirosis.

In this study, we focused on one of the most enigmatic and important aspects of *B. burgdorferi’s* biology: genome inheritance. Faithful genome inheritance during cellular replication is essential for the propagation of all life forms. Despite its small size of ∼1.5 megabases (Mb), the *B. burgdorferi* genome is the most segmented bacterial genome known to date^15^. It is composed of one linear chromosome and over 20 linear or circular plasmids, several of which have essential roles during the spirochete’s natural tick-vertebrate transmission cycle^4,16–18^. Quantitative polymerase chain reaction (qPCR)-based measurements generated the prevailing view that *B. burgdorferi* has about one chromosome copy per cell^19^. Plasmids are present in a ∼1:1 ratio to the chromosome^20–23^. Given the large size of *B. burgdorferi* cells (10 to 25 µm or longer depending on the strain)^24–26^, this presumed monoploidy suggests that the replicated chromosome and plasmids have to segregate over long distances to ensure their faithful inheritance during division. This has, however, not been examined experimentally.

Experiments in a heterologous *E. coli* system, together with a transposon screen in *B. burgdorferi,* have suggested that the replication and partitioning of *B. burgdorferi* plasmids are mediated by specific plasmid-encoded proteins via unknown mechanisms^18,27–29^. On the other hand, chromosome segregation is predicted to involve a ParA/ParB system^17,30^. ParA and ParB proteins are well-known to work together to mediate the segregation of duplicated chromosomal origins of replication (*oriC*) in a broadly diverse bacteria^30,31^. ParB is often referred to as a “centromere-binding” protein because it specifically binds to centromere-like sequences (*parS*) usually located near *oriC*^30^. After loading on the DNA at the *parS* sites, ParB spreads onto adjacent sequences^32–34^ to form a partition complex. ParA is an ATPase that dimerizes upon ATP binding, which in turn promotes the nonspecific binding of the ParA dimer to the DNA (the nucleoid)^35,36^. Upon interaction with ParA, the ParB-rich partition complex stimulates the ATPase activity of ParA, causing dimer dissociation and release of ParA from the DNA^36,37^. Repetition of this biochemical cycle, combined with a translocation force^38^ derived from the elastic properties of the chromosome^39–42^ and/or a diffusion-based mechanism^43–46^, drives the translocation and therefore the segregation of replicated partition complexes^31,47^. The *B. burgdorferi* chromosome contains a *parS* site^30^ near *oriC* and encodes both ParA and ParB homologs^17^, predictive of a conserved ParA/ParB function in chromosome segregation.

In this study, we genetically labeled and imaged various chromosomal loci and plasmids in live *B. burgdorferi* cells. Fluorescence microscopy analysis, combined with genetic deletions and ChIP-seq experiments, revealed that *B. burgdorferi* is polyploid and uses a novel centromere-binding protein, rather than ParB, to carry out a key DNA partitioning activity.

## RESULTS

### B. burgdorferi carries multiple chromosome copies per cell during growth in culture

To label *oriC* in live *B. burgdorferi* cells, we relied on the specific recognition of the *oriC*-proximal *parS* site by ParB (see supplemental text and Extended Data Fig. 1A-B). We therefore substituted the *parB* gene with *mcherry-parB* at the endogenous locus. The resulting strain (CJW_Bb474), which also expressed free GFP for cytoplasm visualization, was stained with the Hoechst dye to reveal DNA localization. In this strain, we expected to detect one or two fluorescent mCherry-ParB puncta within the nucleoid, reflecting a monoploid state before and after initiation of chromosomal DNA replication, respectively. To our surprise, each cell had multiple mCherry-ParB foci that appeared regularly spaced along the nucleoid (Fig. 1A).

**Figure 1.**
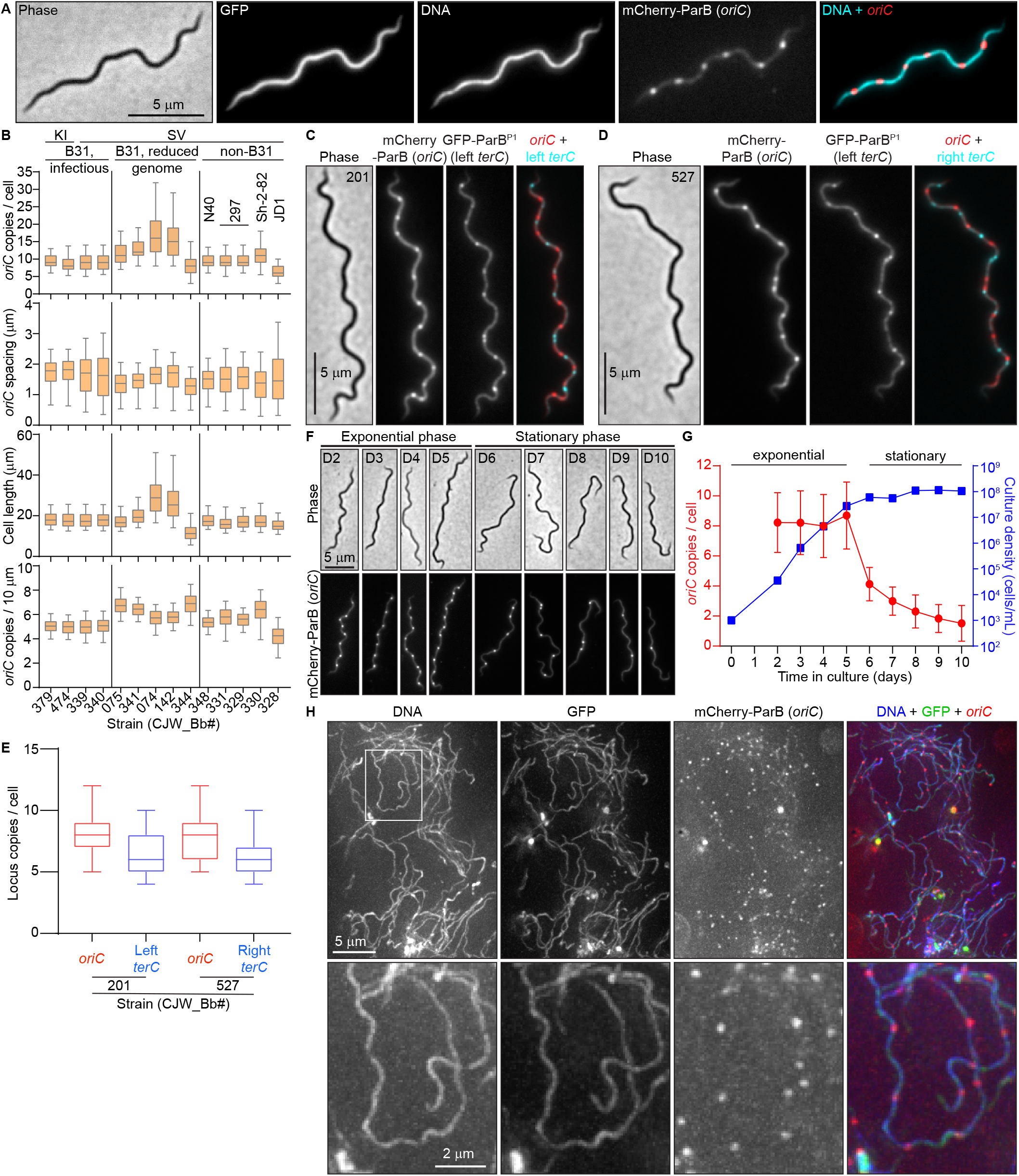
*B. burgdorferi* cells contain multiple chromosome copies. **A.** Images of a cell of strain CJW_Bb474 expressing cytosolic free GFP and mCherry-ParB. Hoechst 33342 was used to stain the DNA. **B.** Population quantifications of exponentially growing cultures for the strains shown in Extended Data Fig. 1C. Top to bottom: Copy numbers per cell of the labeled *oriC* loci, *oriC* spacing (distances between adjacent *oriC* spots); cell lengths, and *oriC* densities (copies per 10 μm of cell length). Strain numbers are indicated at the bottom. Shown are the mean of the data (middle line), the 25 to 75 percentiles of the data (box), and the 2.5 to 97.5 percentiles of the data (whiskers). Strain backgrounds and *oriC* labeling method (*mcherry-parB* knock-in, KI, or tagged *parB* expressed from a shuttle vector, SV) are listed at the top. **C.** Phase-contrast and fluorescence micrographs of a cell of strain CJW_Bb201, which expresses labels for *oriC* (mCherry-ParB) and the left *terC* locus (GFP-ParB^P1^), respectively. **D.** Phase-contrast and fluorescence micrographs of a cell of strain CJW_Bb527, which expresses labels for *oriC* (mCherry-ParB) and the right *terC* locus (GFP-ParB^P1^), respectively. D and E. Images were acquired while the cultures were growing exponentially. **E.** Boxplots of *oriC* and *terC* copies per cell in exponentially growing cultures of strains CJW_Bb201 and CJW_Bb527. Shown are the mean of the data (middle line), the 25 to 75 percentiles of the data (box) and the 2.5 to 97.5 percentiles of the data (whiskers). **F.** An exponentially growing culture of strain CJW_Bb339, in which *oriC* is labeled by expression of mCherry-ParB, was diluted to 10^3^ cells/mL then imaged daily from day 2 through day 10 of growth in culture. Shown are representative images of cells from the days indicated on the phase-contrast images. **G.** Plot showing the *oriC* copy number per cell (red, mean ± standard deviation) and the culture density (blue, in cells/mL) over time for the population imaged in (F). **H.** Images of cells of strain CJW_Bb474 in the midgut of a nymphal tick 10 days after feeding drop-off. DNA was stained with Hoechst 33342. The mCherry-ParB *oriC* signal was amplified by staining with RFP booster. Shown are max-Z projections of a deconvolved stack of images (top) and higher-magnification views of the region indicated by the white rectangle.

We confirmed this finding in 13 additional *B. burgdorferi* strains derived from different lineages of the strain B31, or from other *B. burgdorferi* isolates, including the well-studied N40 and 297 strains (Fig. 1B and Extended Data Fig. 1C; also see supplemental text for strain construction and *oriC* labeling). The spacing between adjacent *oriC* copies was similar across all tested strains (Fig. 1B). As a complementary approach, we labeled the *oriC*-proximal *uvrC* locus of the *B. burgdorferi* B31 chromosome with the orthogonal GFP-ParB^P1^/*parS^P1^* pair derived from the *E. coli* P1 plasmid and observed similar results (see supplemental text and Extended Data Fig. 1A-B, D-F). To demonstrate that the detection of multiple *oriC* copies reflects the presence of multiple chromosomes per cell, we genetically labeled and visualized the left and right telomeres (*terC*) using the GFP-ParB^P1^/*parS^P1^* system (Fig. 1C-E). We also confirmed the presence of multiple *terC* copies per cell by DNA fluorescence *in situ* hybridization (Extended Data Fig. 1G). Collectively, our data show that *B. burgdorferi* cells contain multiple complete chromosomes and thus are polyploid during exponential growth in culture.

We found that the discrepancy between our polyploidy results and the qPCR measurements of ∼1.3 chromosomes per cell^19^ stemmed from a difference in culture growth stage. While we imaged cultures in exponential growth (Fig. 1A-E, Extended Data Fig. 1B-G), the previous analysis was done using “saturated” cultures^19^, which had likely reached stationary phase. Indeed, we found that the *oriC* copy number decreases in stationary phase cultures, ultimately reaching about one copy per cell (Fig. 1F-G).

### B. burgdorferi cells contain multiple copies of their endogenous plasmids

Next, we examined the localization and copy number of 16 distinct plasmids relative to the chromosome by generating different strains, each with a distinct endogenous plasmid labeled in addition to the *oriC* locus (Fig. 2A, Table S1). For each plasmid, we detected multiple copies per cell (Fig. 2A, Extended Data Fig. 2A). The ratio of plasmid and *oriC* copies varied between 0.5 and 2 (Fig. 2B, Extended Data Fig. 2A), in agreement with previous findings^20–22^. These results were further confirmed by marker frequency analysis (Fig. 2B). Moreover, the plasmids were regularly spaced within the cells (Fig. 2A). As with the chromosome, the copy number of plasmids (e.g., cp26) decreased as the *in vitro* culture advanced into stationary phase (Extended Fig. 2B), suggesting a potential coordination between independent segments of the genome.

**Figure 2.**
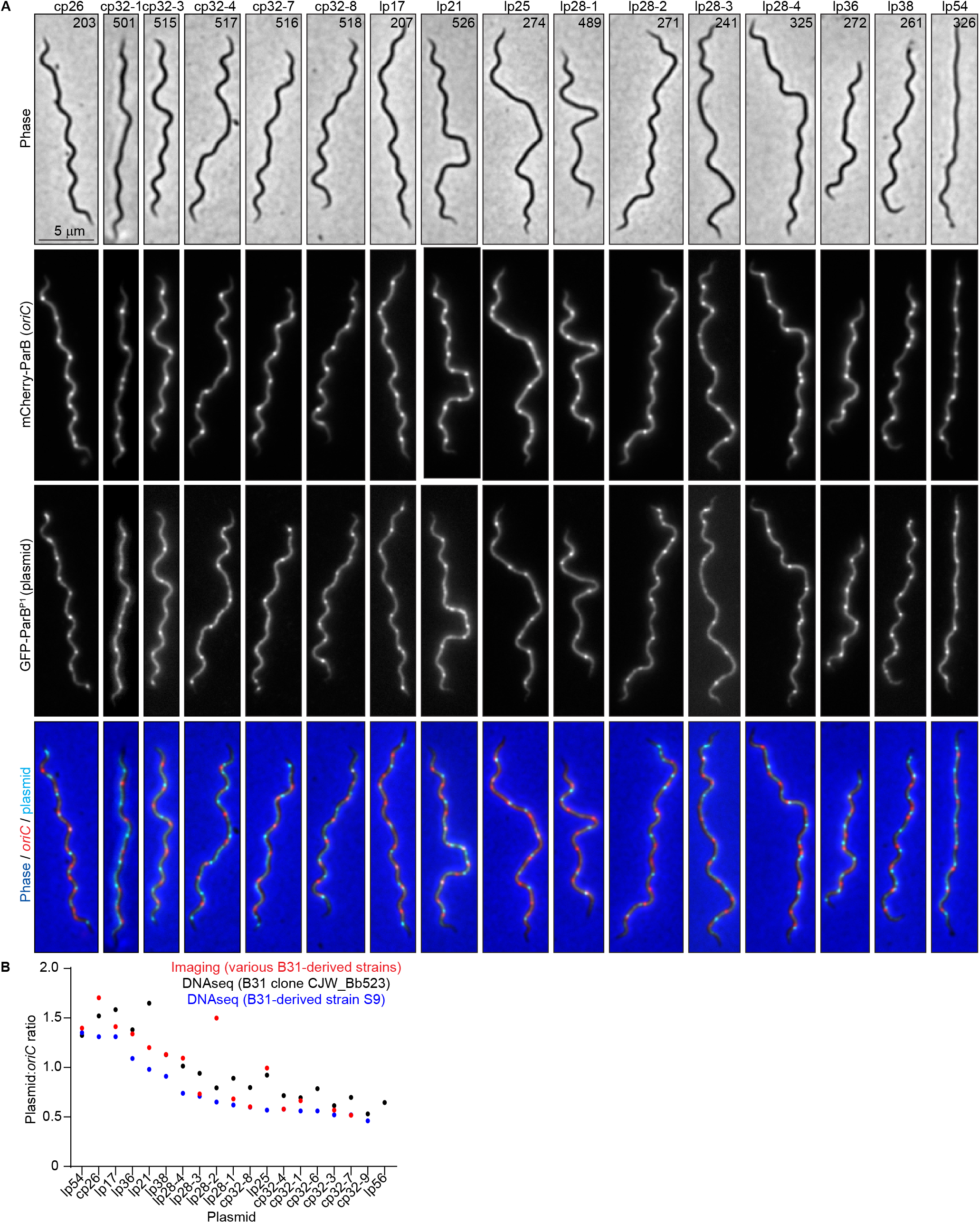
Exponentially-growing *B. burgdorferi* cultures contain multiple copies per cell of\ their endogenous plasmids. **A.** Images of representative cells of strains in which the *oriC* loci were labeled by expression of mCherry-ParB while the plasmid indicated at the top was labeled by insertion of a *parS^P1^* sequence and expression of GFP-ParB^P1^. CJW_Bb strain numbers are marked on the phase-contrast images. **B.** Plot showing the plasmid-to-*oriC* copy number ratios determined by imaging the strains shown in (A) (red), or by DNAseq (see methods) in strains S9 (blue) or a clone of strain B31MI (black).

### B. burgdorferi cells in fed nymphs are also polyploid

While the phenotypes of stationary phase cultures were interesting (and will be explored in a separate study), we chose to focus our attention on *B. burgdorferi* during growth because cell proliferation is required for disease to occur. Outside the laboratory, *B. burgdorferi* cannot grow as a free-living organism. Therefore, to test whether the polyploidy observed in growing cultures has physiological relevance in a natural context, ticks were colonized with strain CJW_Bb474 expressing mCherry-ParB and cytoplasmic GFP through feeding on infected mice. This strain displayed no apparent defect in mouse infectivity either by needle injection or tick bite, or in acquisition by ticks feeding on infected mice (Extended Data Fig. 1H-J and supplemental text). Using three-dimensional deconvolution of image stacks, we were able to readily detect GFP-positive spirochetes in the midgut of fed, CJW_Bb474-colonized nymphs (Fig. 1H). These spirochetes, which were co-stained with the DNA dye Hoechst, contained regularly spaced mCherry-ParB foci (Fig. 1H). The complex three-dimensional orientation of the thin spirochetes within the tick midgut^48^ prevented us from accurately determining the number of *oriC* copies per cell or measuring distances between adjacent *oriC* copies. Nevertheless, based on the inset in Fig. 1H, we estimated the inter-origin distance to be ∼ 2 μm, which is only slightly larger than the distances measured in growing cultures (Fig. 1B). Importantly, we establish that the polyploidy of *B. burgdorferi* cells is physiologically relevant.

### B. burgdorferi oriC copy numbers correlate with cell length

In culture, strains with longer cells had higher *oriC* copy numbers per cell (Fig. 1B). Correlation between *oriC* copy number and cell length was also apparent at the single-cell level for each strain (Extended Data Fig. 3A-B), a scaling property that persisted after blocking cell division using the FtsI inhibitor piperacillin^49^ (Fig. 3A-B). *terC* copy numbers also correlated linearly with cell length (Fig. 3C-D), as did plasmid copy numbers (Extended Data Fig. 2C-D). We therefore approximated chromosome and plasmid densities by calculating the number of copies found in 10 μm of cell length (Fig. 1B and Extended Data Fig. 2A). Among the B31-derived strains, those with reduced genomes (i.e., having fewer endogenous plasmids) had higher *oriC* densities than those with more endogenous plasmids (Fig. 1B), suggesting that *B. burgdorferi* may initiate DNA replication in response to the cellular space available to be filled by the DNA.

**Figure 3.**
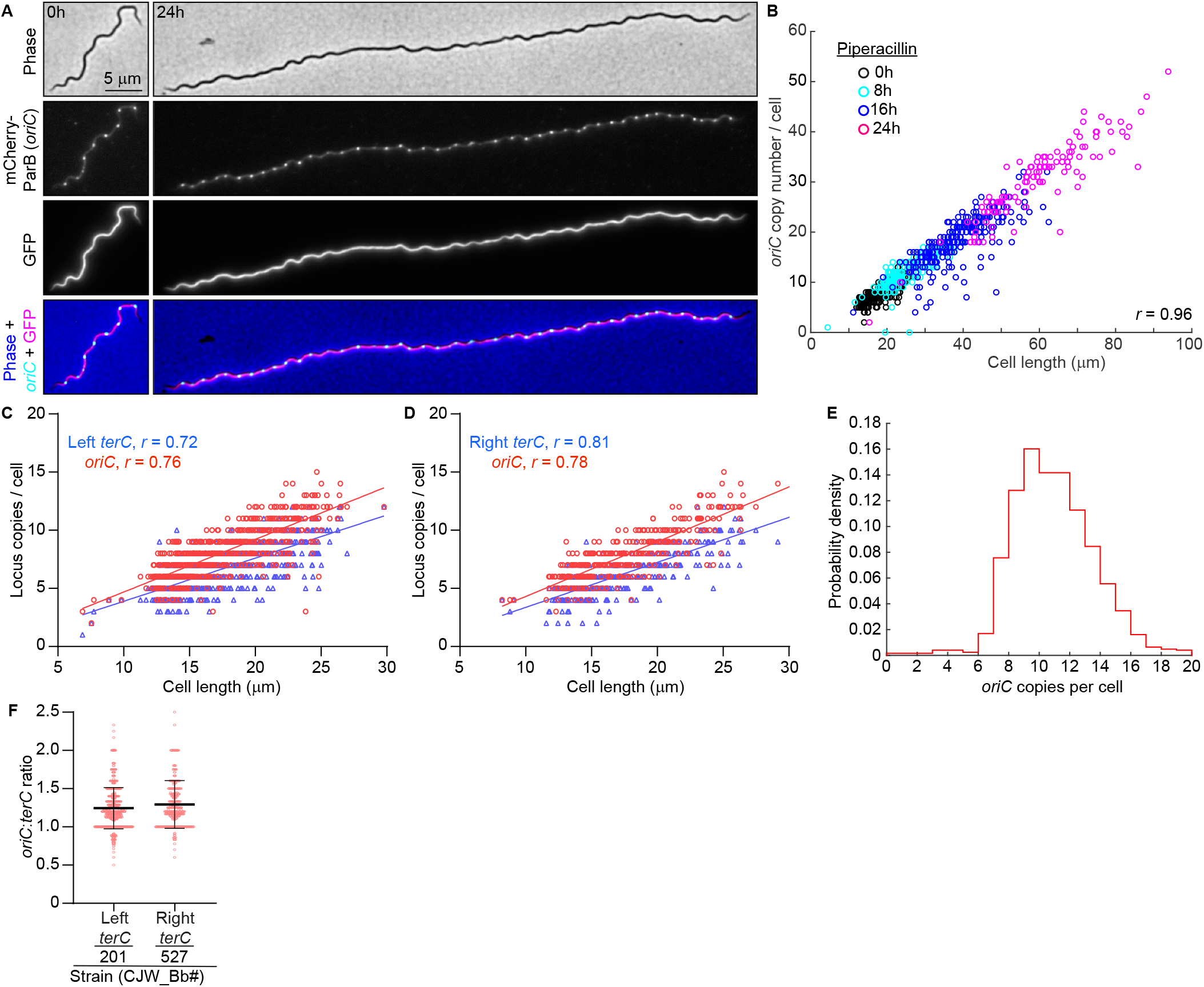
*B. burgdorferi* chromosome copy numbers correlate with cell length. **A.** Images of cells of strain CJW_Bb474 before (0 h) or after 24 h of exposure to piperacillin. **B.** Correlation of *oriC* copy number per cell with cell length in strains CJW_Bb474 and CJW_Bb379 following the indicated durations of piperacillin treatment. *r*, Pearson’s correlation coefficient. **C.** Correlations of *oriC* and *terC* copy number per cell with cell length in exponentially growing cultures of strain CJW_Bb201. *r*, Pearson’s correlation coefficient. **D.** Correlations of *oriC* and *terC* copy number per cell with cell length in exponentially growing cultures of strain CJW_Bb527. *r*, Pearson’s correlation coefficient. **E.** Histogram of *oriC* copy numbers per cell in exponentially growing cultures of strain CJW_Bb379. **F.** Plot showing the *oriC*:*terC* ratios calculated in single cells from exponentially growing cultures of strains CJW_Bb201 and CJW_Bb527, following fluorescent spot detection. Black lines depict means ± standard deviations.

Several lines of evidence suggest that *B. burgdorferi* initiates DNA replication asynchronously. First, the *oriC* copy number in exponentially growing *B. burgdorferi* cultures had a unimodal distribution (Fig. 3E), as opposed to the expected bimodal distribution if all *oriC* copies would replicate at the same time. Second, at the population level, the *oriC*:*terC* ratio was ∼1.2 on average, as measured both by imaging and by marker frequency analysis using whole genome sequencing (see methods). Such an average *oriC*:*terC* ratio close to 1 indicates that few chromosomes are replicating at the same time, which we also confirmed at the single-cell level, as most individual cells had *oriC*:*terC* ratios smaller than 2 (Fig. 3F). Asynchronous chromosome replication and scaling between genome copy and cell length may be a common property of polyploid bacteria, as they are also observed in cyanobacteria^50,51^.

### B. burgdorferi ParA is depleted at oriC loci

We next investigated the regular spacing of chromosome copies along the cell length, which ensures near-even distribution of the chromosome to daughter cells following division at midcell. Based on the known functions of ParA and ParB in other bacteria^31,47,52^, we suspected that the *B. burgdorferi parA* and *parB* genes, which reside close to the predicted *parS* site (Fig. 4A), played an important role in the regular patterning of *oriC* copies in *B. burgdorferi*. As expected, we found that the formation of mCherry-ParB foci was dependent on the presence of *parS* (Extended Data Fig. 1B and supplemental text). The genome-wide binding profile of mCherry-ParB, determined by ChIP-seq, revealed a single enrichment peak at the *parS* site (Fig. 4A-B; also see Extended Data Fig. 6B and supplemental text). Furthermore, chromosomal replacement of *parA* with *parA-msgfp* revealed an uneven ParA-GFP signal distribution along the length of the cell (Fig. 4C). The regions of concentrated ParA-GFP signal alternated with regions of signal depletion that corresponded to the location of the mCherry-ParB foci in these cells, creating a banded localization pattern (Fig. 4C-E). While ParA-GFP displayed modest accumulation between mCherry-ParB foci that were in close proximity, it accumulated prominently between mCherry-ParB foci that were farther apart (Fig. 4D-E). This localization pattern is expected for ParA/ParB DNA partitioning systems^40,41^, as ParB is known to stimulate ParA depletion^53,54^.

**Figure 4.**
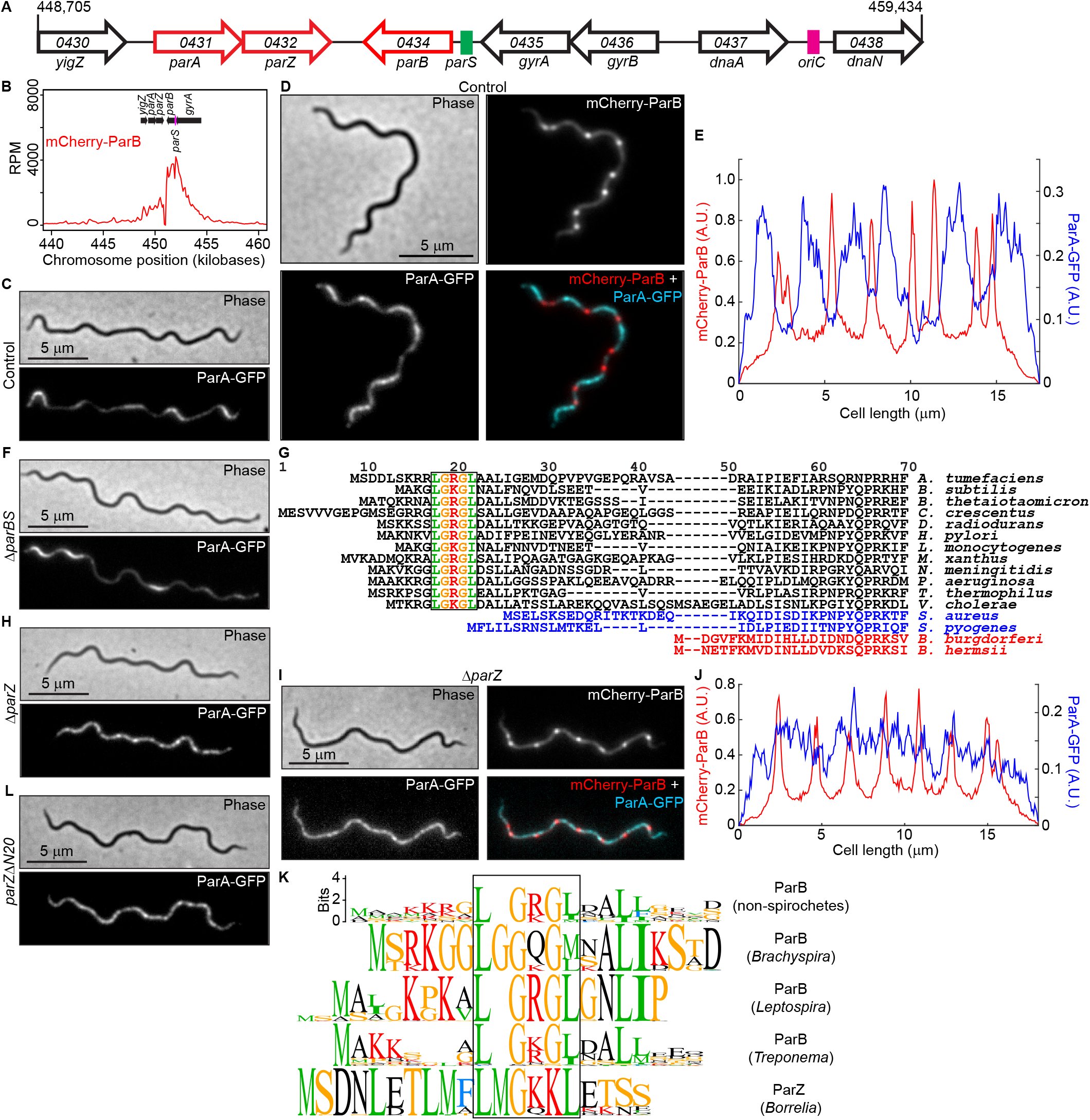
ParZ, not ParB, controls ParA localization. **A.** Schematic of the *oriC*-proximal chromosome region of *B. burgdorferi* strain B31, between nucleotides 448,705 and 459,434, showing the *par* genes. Features are not drawn to scale. We propose renaming the gene *bb0432* of hypothetical function to *parZ*. **B.** ChIP-seq profile showing the binding of mCherry-ParB on the chromosome in strain CJW_Bb379. Only the genome coordinates corresponding to the *par* locus and surrounding DNA sequences are shown (see Extended Data Fig. 6B for the whole genome). Genes underlining the enriched sequence reads are highlighted. RPM, reads per million. **C.** Images of ParA-GFP localization in knock-in strain CJW_Bb488, which carries a full complement of *par* genes and thus serves as control for ParA-GFP localization. **D.** Images of a cell of strain CJW_Bb538 expressing ParA-GFP and in which *oriC* copies were localized by expression of mCherry-ParB. **E.** Fluorescence intensity profiles along the cell length for the cell shown in (D). **F.** Images of ParA-GFP localization in knock-in strain CJW_Bb520 in which *parBS* is deleted. **G.** Alignment of chromosomally encoded ParB sequences from the indicated species. The first 70 positions of the alignment are shown here, while the full alignment is displayed in Extended Data Fig. 4A. Highlighted is the conserved LG(R/K)GL motif of ParB’s N-terminal ParA ATPase-stimulating peptide, which is absent from *Borrelia, Streptococcus,* and *Staphylococcus* ParB sequences. **H.** Images of ParA-GFP localization in knock-in strain CJW_Bb519 in which *parZ* is deleted. **I.** Images of a cell of strain CJW_Bb619 expressing ParA-GFP and in which *parZ* is deleted and *oriC* copies are localized by expression of mCherry-ParB. **J.** Fluorescence intensity profiles along the cell length for the cell shown in (I). **K.** Sequence logos for the N-terminal 20 amino acids obtained by ClustalW alignment of unique *Borrelia* ParZ sequences as well as unique ParB sequences from selected non-spirochete bacteria (see panel G) or from spirochete *Leptospira*, *Brachyspira*, or *Treponema* genera. Only chromosome I sequences were considered for *Leptospira*. Boxed in are motifs similar to the conserved motif found within the N-terminal peptide of the more diverse chromosomal ParB sequences shown in (G). **L.** Images of ParA-GFP localization in knock-in strain CJW_Bb610 in which *parZ* is missing the sequence encoding the N-terminal 20 amino acids (*parZ*Δ*N20*).

### ParZ, not ParB, controls ParA localization in B. burgdorferi

As ParB normally controls ParA localization in other bacterial ParA/ParB systems^55,56^, we anticipated that deletion of *parB* and *parS* (*parBS*) would disrupt the ParA banded pattern and result in a more uniform distribution. Surprisingly, deleting *parBS* did not eliminate the banded ParA-GFP localization in *B. burgdorferi* (Fig. 4F). Seeking an explanation for this phenotype, we inspected the *B. burgdorferi par* system more closely. We noticed that *B. burgdorferi* ParB lacked an otherwise conserved N-terminal peptide (Fig. 4G, Extended Data Fig. 4A), which is required for stimulation of ParA ATPase activity in other ParA/ParB systems^36,37^. The absence of this peptide from *B. burgdorferi* ParB explains its inability to control ParA localization. The ParB proteins of staphylococci and streptococci also lacked this N-terminal peptide (Fig. 4G, Extended Data Fig. 4A). However, these bacteria also lack a ParA homolog^30,57,58^ and thus do not need the ParA control function of ParB. The rest of the *B. burgdorferi* ParB sequence is similar to that of other chromosomal ParB proteins (Extended Data Fig. 4A). *B. burgdorferi* ParA also has a typical chromosomal ParA sequence (Extended Data Fig. 4B).

Since ParB did not control ParA localization in *B. burgdorferi*, we hypothesized that another factor fulfilled this function. Interestingly, in *B. burgdorferi*, the *parA* and *parB* genes are located on the chromosome in opposite, head-to-head orientations (Fig. 4A), instead of being joined in a two-gene operon as in many other bacteria^59^. This opposing gene organization is unique to Borreliaceae as *parA* and *parB* are organized in an operon structure in other spirochetes (Extended Data Fig. 4C). Furthermore, *parA* appears to form an operon with *bb0432*, a gene of hypothetical function that we propose to rename *parZ* (Fig. 4A). We identified putative *parAZ* operons among the 29 sequenced Lyme disease *Borrelia* strains that we examined (Extended Data Fig. 4D). ParZ proteins are also well conserved across the entire sequence among the larger Borreliaceae family, including relapsing fever spirochetes (Extended Data Fig. 4E), but are not encoded by other spirochete genomes. Importantly, deletion of *parZ* in *B. burgdorferi* drastically altered the distribution of the ParA-GFP signal (Fig. 4H): the banded pattern of ParA-GFP disappeared and the fluorescent signal became more distributed within the cell, forming only patches, likely due to the known cooperativity of ParA binding to the DNA^35,60^. Additionally, in a Δ*parZ* background, we did not detect depletion of the ParA signal from the vicinity of *oriC*, nor did we see banded accumulation of ParA-GFP in cellular regions located between the *oriC* loci (Fig. 4I-J). Thus, ParZ regulates the subcellular localization of ParA in *B. burgdorferi*.

In many bacteria, ParB controls ParA activity via a basic catalytic residue^36^ that lies within a conserved motif, LG(R/K)GL, located in the N-terminal peptide (Fig. 4G). Related motifs can also be found within the N-terminal peptides of chromosomal ParB sequences from spirochetes outside Borreliaceae (Fig. 4K). Intriguingly, ParZ sequences had a similar motif in their well-conserved N-terminal peptide (Fig. 4K and Extended Data Fig 4E). Removal of this N-terminal ParZ peptide (ParZΔN20) was sufficient to disrupt ParA-GFP localization (Fig. 4L), even though the peptide deletion had no apparent effect on transcription (Extended Data Fig. 5). Collectively, our data indicate that ParZ substitutes ParB’s function in controlling ParA localization using a similar N-terminal motif.

### ParZ is a novel bacterial centromere-binding protein

If ParZ substitutes ParB in ParA-mediated chromosome segregation, we reasoned that ParZ must directly or indirectly bind DNA close to *oriC* to explain the ParA depletion at mCherry-ParB foci (Fig. 4D-E). Indeed, chromosomal replacement of *parZ* with *parZ-msfgfp* revealed regularly spaced fluorescent foci (Fig. 5A, WT) that resembled *oriC* labeling by mCherry-ParB (Figs. 1A and 5B). The copy number and density of the ParZ-GFP foci were similar to those of the mCherry-ParB foci (Extended Data Fig. 6A, compare the two control strains) and the ParZ-GFP and mCherry-ParB foci colocalized (Fig. 5C-D). As ParZ-GFP formed foci in the absence of *parBS* or *parA* (Fig. 5A), ParZ localization at the *oriC* regions suggested the presence of a novel centromere-like region near *oriC*. Indeed, ChIP-seq experiments using ParZ-GFP identified a specific enrichment region that centers at the *parAZ* region (Fig. 5E). Strikingly, this enrichment peak was adjacent to but distinct from the mCherry-ParB ChIP-seq peak centered on the *parS* site (Fig. 5E, Extended Data Fig. 6B-C and supplemental text). Importantly, the ParZ-GFP ChIP-seq peak was preserved in the Δ*parBS* background (Fig. 5F), as was the mCherry-ParB peak in the Δ*parAZ* background (Fig. 5G), in agreement with our microscopy observations (Fig. 5A-B). Both proteins spread over a total of ∼8 kb-DNA sequences upstream and downstream of their encoding genes (Fig. 5E and Extended Data Fig. 6C). To test whether the ParZ binding site is within the *parZ* gene, we performed ChIP-seq in a strain that has a copy of *parZ-msgfp* on a shuttle vector. We found that having the *parZ* gene sequence on the shuttle vector was sufficient to recruit ParZ-GFP to the vector (Extended Data Fig. 6D-G and supplemental text). Thus, ParZ binds within its own gene.

**Figure 5.**
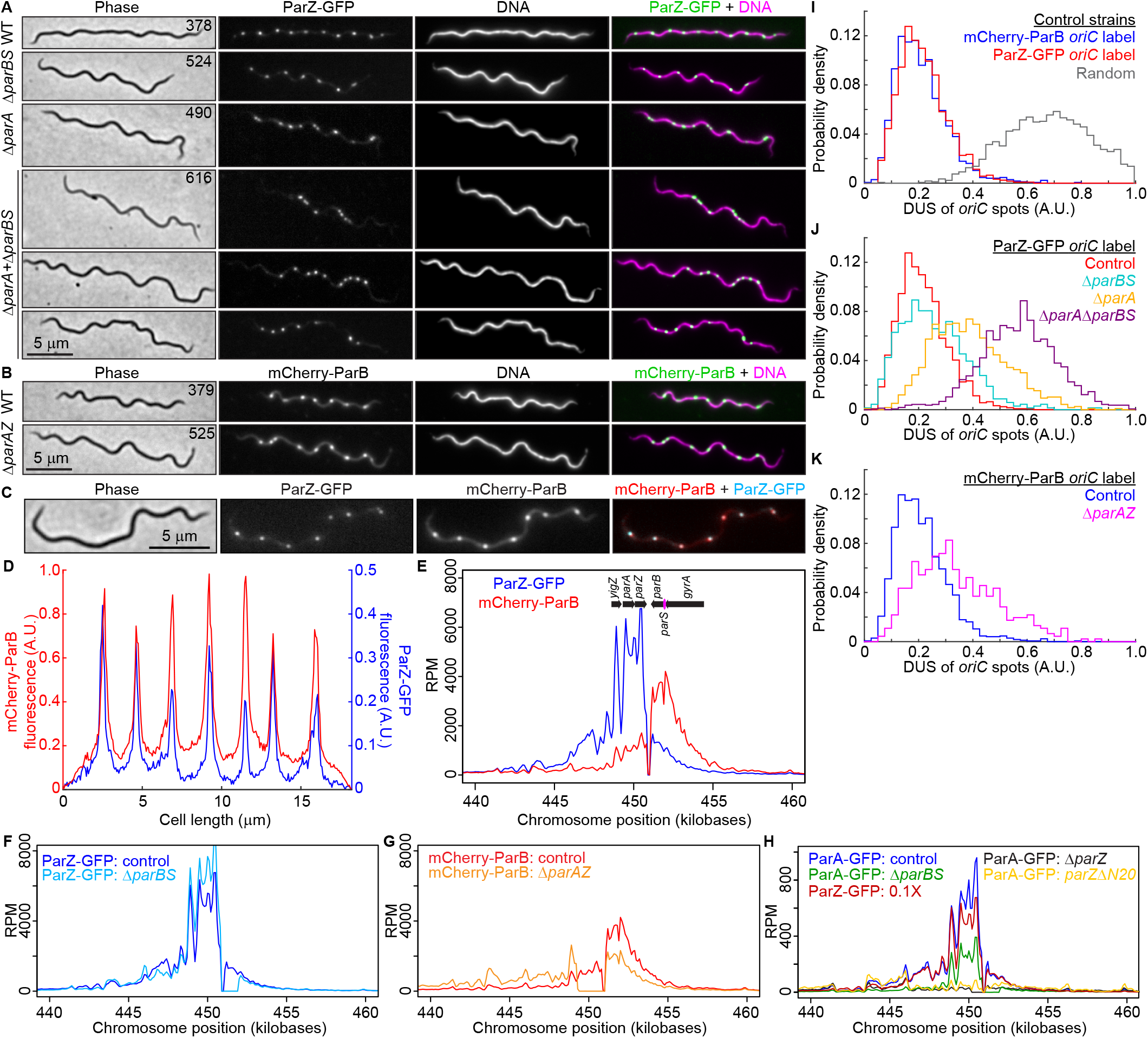
ParZ is a novel bacterial centromere-binding protein that contributes to *oriC* segregation. **A.** Images of ParZ-GFP signal in strains carrying the full complement of *B. burgdorferi par* genes (WT) or lacking the *par* genes and/or *parS* sequence as indicated. DNA was stained with Hoechst 33342. CJW_Bb strain numbers are listed on the phase-contrast images. **B.** Images of mCherry-ParB signal in strains carrying the full complement of par genes (WT) or lacking the *parAZ* operon, as indicated. DNA was stained with Hoechst 33342. CJW_Bb strain numbers are listed on the phase contrast images. **C.** Images of strain CJW_Bb403 showing colocalization of ParZ-GFP and mCherry-ParB signals. **D.** Fluorescence intensity profile along the cell length for the cell shown in (C). **E.** ChIP-seq profiles showing the binding of ParZ-GFP and mCherry-ParB on the chromosome using strains CJW_Bb378 and CJW_Bb379, respectively. The mCherry-ParB trace is the same as the one shown in Fig. 4B. Only the genome regions corresponding to the *par* locus and surrounding DNA sequences are shown (see Extended Data Fig. 6B for the whole genome). Genes underlining the enriched sequence reads are highlighted. RPM, reads per million. The dip in the trace between *parZ* and *parB* is due to read mapping to the reference B31 chromosomal sequence which does not contain the kanamycin resistance cassette present in the strains analyzed by ChIP-seq (see supplemental text). The same is true for (F-H). **F.** ChIP-seq profiles comparing binding of ParZ-GFP to the *par* locus in the presence (control, strain CJW_Bb378) or absence of *parBS* (strain CJW_Bb524). **G.** ChIP-seq profiles comparing binding of mCherry-ParB to the *par* locus in the presence (control, strain CJW_Bb379) or absence of *parAZ* (strain CJW_Bb525). **H.** ChIP-seq profiles comparing binding of ParA-GFP to the *par* locus in the presence (control, strain CJW_Bb488) or absence of *parBS* (strain CJW_Bb520) or *parZ* (strain CJW_Bb519). Also shown is ParA-GFP binding to the *par* locus in the presence of *parZ*Δ*N20* (strain CJW_Bb610). Overlayed is a scaled-down (0.1 X) ParZ-GFP ChIP-seq profile in the control background (strain CJW_Bb378, see blue trace in panel E). **I.** Histograms showing the distributions of deviations from uniform spacing (DUS, see methods) of *oriC* spots in control strains expressing mCherry-ParB (strain CJW_Bb379) or ParZ-GFP (strain CJW_Bb378) to label *oriC*. Also shown in gray are DUS values obtained by simulating a random redistribution of *oriC* loci in the analyzed cells of strain CJW_Bb379 (see methods). **J.** Histograms comparing DUS distributions of ParZ-GFP-labeled *oriC* foci in control (CJW_Bb378), Δ*parBS* (CJW_Bb524), Δ*parA* (CJW_Bb490), and Δ*parA* Δ*parBS* (CJW_Bb616) strains. **K.** Histograms comparing DUS distributions of mCherry-ParB-labeled *oriC* foci in control (CJW_Bb379) and Δ*parAZ* (CJW_Bb525) strains.

**Figure 6.**
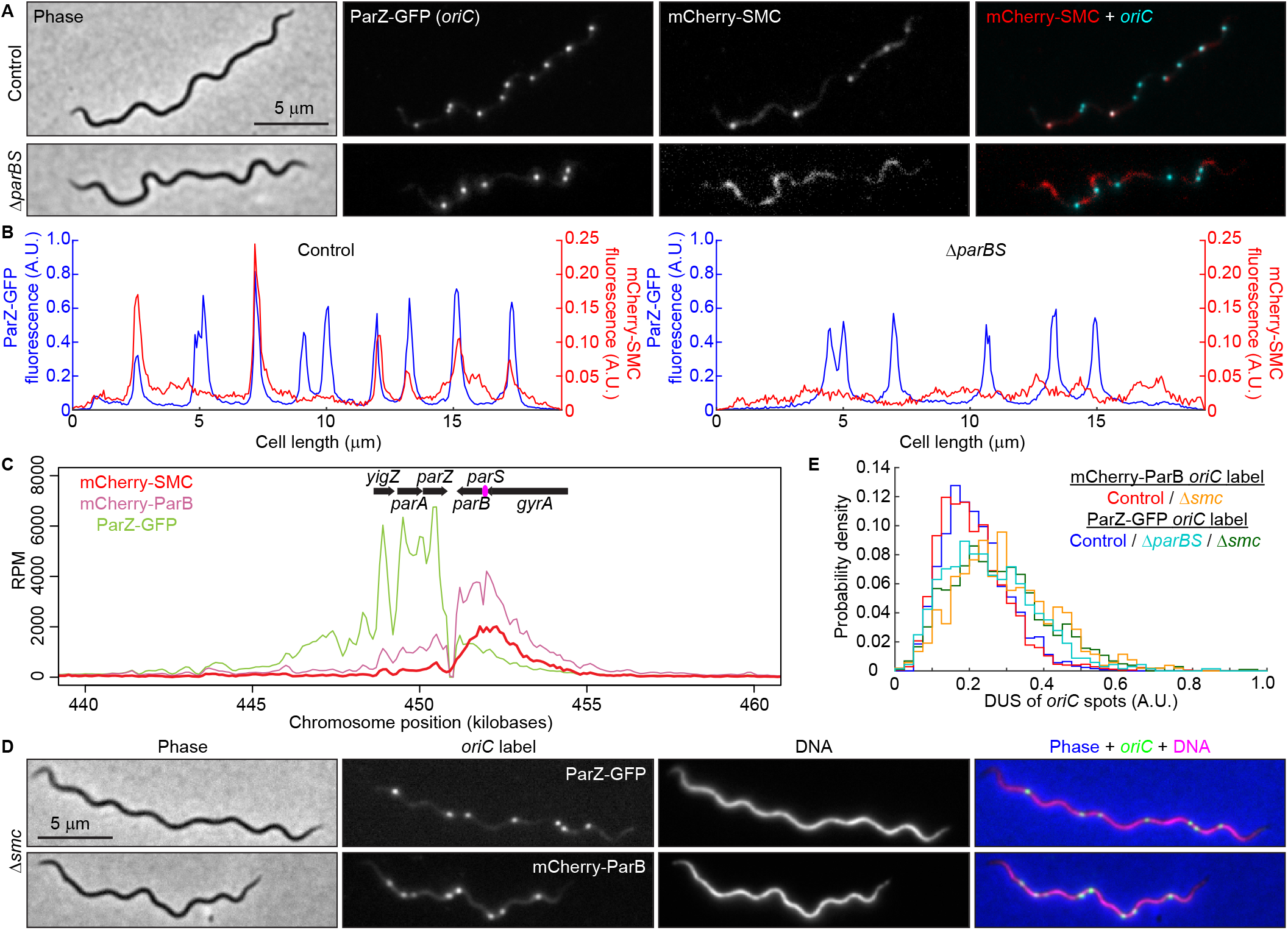
SMC is recruited to the *oriC* region by ParB and mildly contributes to regular *oriC* spacing. **A.** Images of cells expressing ParZ-GFP (to label *oriC*) and mCherry-SMC in control (CJW_Bb602, carrying the full complement of *par* genes) and Δ*parBS* (CJW_Bb615) strains. **B.** Fluorescence intensity profiles along the cell length for the cells shown in (A). **C.** ChIP-seq profiles comparing binding of mCherry-SMC (red, strain CJW_Bb601), ParZ-GFP (green, strain CJW_Bb378) and mCherry-ParB (pink, strain CJW_Bb379). Genes underlining the enriched sequence reads are highlighted. RPM, reads per million. The dip in the trace between *parZ* and *parB* is due to read mapping to the reference B31 chromosomal sequence which does not contain the kanamycin resistance cassette present in the strains analyzed by ChIP-seq (see supplemental text). **D.** Images of cells showing *oriC* localization in Δ*smc* strains in which the *oriC* is labeled with either ParZ-GFP (CJW_Bb603) or mCherry-ParB (CJW_Bb604). Hoechst 33342 was used to stain the DNA. An overlay is also shown. **E.** Histograms showing DUS of *oriC* spots in cells of control and Δ*smc* strains in which oriC was labeled with either ParZ-GFP (control, CJW_Bb378; Δ*smc*, CJW_Bb603) or mCherry-ParB (control, CJW_Bb379; Δ*smc*, CJW_Bb604). Also shown is the DUS histogram for the ΔparBS strain CJW_Bb524.

In ParA/ParB systems, ParB bound to *parS* and adjacent sequences controls ParA localization by transiently interacting with ParA^31^. If ParZ works via a similar mechanism in *B. burgdorferi*, it predicts a transient interaction between ParZ and ParA. ChIP-seq experiments verified this prediction as the ParA-GFP enrichment peak is at the same location as the ParZ-GFP peak, albeit at a lower level consistent with a transient interaction (Fig. 5H and Extended Data Fig. 6B,H,I). Furthermore, this ParA-GFP binding did not require *parBS* but required ParZ and, more specifically, its N-terminal peptide (Fig. 5H). These results indicate that ParZ is a newly identified centromeric protein that, similarly to ParB in other bacteria, uses its N-terminal peptide to regulate ParA’s localization.

### ParA, ParB, and ParZ jointly control chromosomal oriC spacing

Given that ParZ, and not ParB, controls ParA in *B. burgdorferi*, we investigated the role of each Par protein in *oriC* segregation using a comprehensive set of *par* gene deletion mutants. In the absence of an appropriate setup for live-cell timelapse imaging of *B. burgdorferi*, we analyzed static snapshots of exponentially growing populations of cells. Visual inspection of the control strains that express mCherry-ParB or ParZ-GFP to label *oriC* revealed a near-equidistant spacing of *oriC* copies along the cell length (Fig. 5A-B, WT). For quantification, we calculated how much the *ori*C distribution in each cell of a population deviates from uniform spacing (see methods). This metric, referred to as “<u>d</u>eviation from <u>u</u>niform <u>s</u>pacing” (or DUS), gives a value of 0 when a cell displays a perfectly equidistant distribution of *oriC* copies. In contrast, a random intracellular distribution of *oriC* copies yields an average DUS value of 0.7 (Fig. 5I and see methods). Control strains, which have a complete set of *parA, parZ, and parB* genes, had very similar DUS distributions centered around ∼0.2 (Fig. 5I), indicating near-equidistant spacing. Deletion of *parBS* reduced the number of *oriC* foci (Extended Data Fig. 6A) but had little impact on the regularity of *oriC* spacing (Fig. 5A). This was reflected in the DUS distribution, which displayed only a slight shift towards larger values relative to the control strains (Fig. 5J). In contrast, deletion of *parA* or of the *parAZ* operon visibly disrupted the spacing of the *oriC* foci (Fig. 5A-B). This was reflected in larger, rightward shifts of DUS distributions (Fig. 5J-K). Deletion of both *parA* and *parBS* caused the strongest *oriC* spacing defect, which manifested as multiple *oriC* foci clustering in one or several short cell segments flanked by large cellular spaces containing DNA devoid of *oriC* spots (Fig 5A). This striking phenotype caused the largest DUS distribution shift towards higher values (Fig. 5J). Furthermore, the severe *oriC* segregation defect seen in the Δ*parA*Δ*parBS* double mutant was accompanied by a large increase in the frequency of cells that lacked *oriC* foci, from <0.3% in control and single-mutant strains to 3.5% in the Δ*parA*Δ*parBS* double mutant. Collectively, our findings indicate that *parA/parZ* and *parB/parS* jointly control *oriC* segregation in *B. burgdorferi*, with *parA/parZ* playing a more prominent role.

### ParB recruits SMC to the chromosomal replication origins

Despite having lost its ability to control ParA, *B. burgdorferi* ParB still participates in *oriC* segregation. Therefore, we investigated whether ParB achieves this function via interaction with another protein. In other bacteria, ParB recruits structural maintenance of the chromosome (SMC) complexes to the *oriC* region^61–63^. The recruitment of SMC to *oriC* and subsequent organization of the *oriC* region have been shown to facilitate *oriC* segregation in *Caulobacter crescentus* and *Bacillus subtilis*^61–67^. While lacking the ParA control motif, the *B. burgdorferi* ParB protein has retained a predicted SMC-binding region located within the N-terminal domain of ParB^63^ (Extended Data Fig. 4A). *B. burgdorferi* also encodes a homolog of SMC at locus *bb0045*. We therefore replaced *smc* with an *mcherry-smc* fusion both in the wild-type background and in a strain that expresses ParZ-GFP to visualize the *oriC* region. mCherry-SMC had no detectable effect on *oriC* copy number (Extended Data Fig. 6A) and formed fluorescent foci that colocalized with some, though not all, ParZ-GFP-decorated *oriC* loci (Fig. 6A-B). Consistent with this colocalization, ChIP-seq experiments revealed enrichment of mCherry-SMC at the *par* locus in a pattern similar to that of mCherry-ParB (Fig. 6C). Deletion of *parBS* abrogated the formation of clear mCherry-SMC puncta and their colocalization with ParZ-GFP decorated *oriC* loci (Fig. 6A-B). Finally, the Δ*smc* mutation caused a similar reduction in *oriC* copy number and density as Δ*parBS* (Extended Data Fig. 6A) and mildly disrupted *oriC* spacing (Fig. 6D-E). These results support the idea that ParB recruits SMC near *oriC*, and that this interaction plays a supporting role in *oriC* segregation in *B. burgdorferi*.

## DISCUSSION

Here we demonstrate that the linear *B. burgdorferi* chromosome exists as multiple copies in cultured and tick-borne spirochetes (Fig. 1A-B and 1H, Extended Data Fig. 1C). Plasmids were also present in multiple copies (Fig. 2A, Extended Data Fig. 2A) and in numbers approximating that of the chromosome^20–22^ (Fig. 2B). Our polyploidy demonstration in live cells dispels the common belief that Lyme disease agents such as *B. burgdorferi* differ in ploidy from the relapsing fever agent *Borrelia hermsii*, which was shown to have multiple genome copies per cell using biochemical population-level measurements^23,68^. In fact, polyploidy may reflect a general property of spirochetes, as qPCR experiments have also demonstrated the polyploid nature of the syphilis spirochete *Treponema pallidum*^69^. Polyploidy may be advantageous under stress. For instance, the presence of multiple genome copies has been shown to increase resistance to DNA damage in cyanobacteria^70^, presumably by facilitating DNA recombination and repair. Polyploid cyanobacteria can also recover from infection by lytic phages as long as not all genome copies are degraded prior to phage inactivation^71^. In the context of a pathogen like *B. burgdorferi*, polyploidy may facilitate persistence in the vertebrate host. Modeling studies have predicted that antigenic variation systems facilitate longer survival of pathogens in immunocompetent hosts when they are present in multiple genomic copies than when encoded on a single-copy genome^68^. The plasmid lp28-1 contains an antigen variation system^72^. Its presence in multiple copies (Fig. 2) may further help this spirochete evade the host antibody response. Our polyploidy finding also has practical implications, one being *B. burgdorferi* load assessment in animal tissues by qPCR, which often relies on the assumption that one genome copy represents one bacterial cell. Actual tissue bacterial loads could be ∼10-fold lower.

In addition to polyploidy, we demonstrated the regular subcellular distribution of both chromosome and plasmid copies (Fig. 1A, 2A, and Extended Data Fig. 1C). In the context of symmetric division (i.e., at midcell)^26^, such a regular linear distribution of a polyploid genome ensures that each daughter cell inherits near-equal copies of the genome, as in cyanobacteria^50,51^. Indeed, severe disruption of near-uniform distribution of the *B. burgdorferi* chromosome in the Δ*parA*Δ*parBS* mutant (Fig. 5A,J) was accompanied by chromosome inheritance defects. Regular genome patterning also implies the existence of active segregation mechanisms. For the *B. burgdorferi* plasmids, the conserved plasmid maintenance loci^29^ are likely responsible for their observed subcellular distribution. For bacterial chromosomes, segregation of the *oriC* regions generally involves ParB-dependent control of ParA activity. Unexpectedly, we found that in *B. burgdorferi,* and presumably other *Borrelia* species, ParA works independently of ParB and its localization is instead controlled by a novel centromere-binding protein, which we named ParZ (Fig. 4-5, Extended Data Fig. 4-6). We envision that the ParA/ParZ system harnesses physical or chemical forces within the cell to drive segregation of the *oriC* loci, similarly to how the ParA/ParB system drives patterning of multicopy plasmids^31,40,47^.

While ParB lost its ability to control ParA localization in *B. burgdorferi*, it retained its role in recruiting SMC to *oriC* (Fig. 6), thereby contributing to *oriC* partitioning, though to a lesser degree than the ParA/ParZ system (Fig. 5J). SMC, possibly through the organization of the *oriC* region^66^, also promotes *oriC* segregation. This is evident from the similar *oriC* segregation defects displayed by the Δ*parBS* and Δ*smc* mutants (Fig. 6E). The ParB/SMC system may also be involved in initiation of DNA replication^73^, as both Δ*parBS* and Δ*smc* mutants have decreased *oriC* copy numbers during exponential growth in culture (Extended Data Fig. 6A).

During the evolution of Borreliaceae, ParB retained a SMC loading function but remarkably transferred its ParA-regulating function to a new player. How could it have happened? BLAST searches allowed us to identify distant homologs of ParZ among the Firmicutes and Fusobacteria (Extended Data Fig. 7). The *Streptococcus*, *Staphylococcus*, *Enterococcus*, and *Bacillus* genera are particularly well represented among the Firmicutes hits (Extended Data Fig. 7). These ParZ-like proteins contain only short regions of homology with the Borreliaceae ParZ proteins and do not include the ParB-like N-terminal peptide. More intriguing though is that a subset of these ParZ-like proteins are encoded by phages (Extended Data Fig. 7). It is tantalizing to speculate acquisition of *parZ* from a phage infection of the spirochete ancestor of the Borreliaceae. In fact, *Borrelia* genomes carry extensive evidence of infection by phages, including those related to known streptococcal phages^18^. Phage insertion into the *parB* gene may have fused the sequence encoding the N-terminal, ParA-controlling peptide of ParB to a phage-encoded protein that binds its own DNA sequence. Subsequent genetic drift of the phage-disrupted spirochete genome may have resulted in inversion of the orientation of the *parB* gene and loss of the remaining phage sequence, generating the current locus structure.

While this evolutionary scenario remains speculative, our findings highlight the plasticity and evolvability of microbial genomes, even in the context of fundamental cellular functions such as genome partitioning. Understanding basic biological functions in non-model species is important, particularly in bacterial pathogens where divergent regulatory mechanisms of essential bacterial functions may provide new targets for specific therapeutic intervention.

## ACKNOWLEDGEMENTS

We thank Stanford University’s Cell Sciences Imaging Core Facility (RRID:SCR_017787) for use of its OMX BLAZE microscope, Stanford Medicine’s Animal Histology Services for generating the thin cryosections of OCT-embedded tick midgut samples, Stanford’s Center for Innovation in *In vivo* Imaging (SCi^3^) for use of its cryostat, and Indiana University Center for Genomics and Bioinformatics for deep sequencing. We are grateful to Drs. Brian Stevenson, Linda Bockenstedt, Erol Fikrig, Nikhat Parveen, Melissa Caimano, and Jon Blevins for sharing strains and plasmids, and David Rudner for anti-GFP and anti-mCherry antibodies. C.N.T. was supported in part by an American Heart Association postdoctoral fellowship (award number 18POST33990330). X.W. was supported by National Institutes of Health R01GM141242. P.A.R. and J.W. were supported by the Intramural Research Program of the National Institute of Allergy and Infectious Diseases, National Institutes of Health. C.J.-W. is a Howard Hughes Medical Institute Investigator. The funders had no role in study design, data collection, analysis, and interpretation, or the decision to submit the work for publication.

## AUTHOR CONTRIBUTIONS

C.N.T. and C. J.-W. designed the research. J.W. and P.A.R. performed the tick-mouse transmission studies, tick dissections, and tick tissue preservation. C.N.T. generated plasmids and strains, imaged the samples, analyzed the images, performed phylogenetic analyses, and visualized the data. Y.X. developed the MATLAB image analysis pipeline with input from C.N.T.. X.K., Z.R., and X.W. performed ChIP-seq and WGS analyses and visualization. I.I. performed WGS analyses. M.S. performed DNA FISH. M.R.S. and N.J. generated plasmids and strains. P.A.R. provided reagents. C.N.T., P.A.R., X.W., and C.J.-W. supervised the project and acquired funding. C.N.T. and C.J-W. wrote the paper with input from all authors.

## ONLINE METHODS

### Bacterial strains and growth conditions

The following *Escherichia coli* cloning strains were used to generate, maintain, and grow the various plasmids used in this study: DH5α (Promega), NEB 5-alpha and NEB 5-alpha F’*l^q^* (New England Biolabs), and XL10-Gold (Agilent). The strains were grown on Luria Bertani agar plates incubated at 30°C or in Super Broth (35 g/L bacto-tryptone, 20 g/L yeast extract, 5g/L NaCl, and 6 mM NaOH) liquid medium with shaking at 30°C. Antibiotics were used at the following concentrations: kanamycin at 50 μg/mL, gentamicin at 15 to 20 μg/mL, spectinomycin or streptomycin at 50 μg/mL, rifampin at 25 μg/mL in liquid culture or 50 μg/mL in plates, and hygromycin B at 100 to 200 μg/mL.

*B. burgdorferi* strains and their generation are detailed in Table S1, Worksheet 1. They were grown in complete Barbour-Stoenner-Kelly (BSK)-II liquid medium in humidified incubators at 34°C under 5% CO_2_ atmosphere^6,10,11^. Complete BSK-H media was acquired from Sigma-Aldrich. Tubes contained 6 to 7 mL or 13 to 15 mL culture depending on the tube size used (8-mL volume, Falcon, #352027, or 16-mL volume, Falcon, #352025) and were kept tightly closed. Any larger volume vessels were kept loosely capped in the incubator. BSK-1.5 medium for plating was previously described^12,74,75^. Selection antibiotics were used at the following final concentrations in both liquid cultures and plates: kanamycin at 200 μg/mL^76^, gentamicin at 40 μg/mL^77^, streptomycin at 100 μg/mL^78^, hygromycin B at 300 μg/mL^75,79^ and blasticidin S at 10 μg/mL^79^. Piperacillin was used at a final concentration of 10 ng/mL. Unless otherwise indicated, all cultures were maintained in exponential growth by diluting the cultures into fresh medium before reaching ∼5×10^7^ cells/mL.

Semisolid BSK-agarose plating medium^75^ contained 2 parts of 1.7% agarose solution in water and 3 parts BSK-1.5 medium containing appropriate amounts of selective antibiotics, as needed, to yield in the final plating mix the concentrations listed above. Each plate was seeded with a maximum of 1 mL *B. burgdorferi* culture. The agarose, melted and maintained at 55°C, and the BSK-1.5 medium, briefly pre-equilibrated at 55°C, were mixed and then 25 mL were poured into each *B. burgdorferi*-seeded plate. The plates were swirled gently to mix, allowed to solidify at room temperature (RT) in a biosafety cabinet for ∼ 30 min, then transferred to a humidified CO_2_ incubator and incubated between 10 days and 3 weeks.

### Genetic manipulations

All plasmids (Table S1, Worksheet 2) were generated using standard molecular biology techniques that included ligation of restriction endonuclease-digested plasmids and PCR products, Gibson assembly^80^ of DpnI-digested PCR products using New England Biolabs’ platform, or site-directed mutagenesis using the Agilent Quick Change Lightning Site-Directed Mutagenesis kit. Sequences of the oligonucleotide primers used in the course of plasmid generation are provided in Table S1, Worksheet 3. The plasmids, for which relevant sequences were confirmed by Sanger DNA sequencing at Quintara Biosciences, were introduced into *E. coli* host strains by heat shock or electroporation and were archived as *E. coli* strains. Minipreps were done with Zymo Research Zippy plasmid miniprep kit. When requesting any plasmid, we urge the requestor to provide us with the *E. coli* CJW strain number listed in Table S1, Worksheet 3, in addition to the name of the plasmid.

Large amounts of plasmid DNA were isolated from saturated 50 mL Super Broth *E. coli* cultures using the Qiagen Plasmid Plus Midi kit with the final elution performed in water. *E. coli* / *B. burgdorferi* shuttle vectors (25 to 50 μg) were electroporated into 50-100 μL volumes of *B. burgdorferi* competent cells, which were prepared as previously described^81^. Suicide vectors (50 to 75 μg) were linearized with the restriction endonucleases indicated in Table S1, Worksheet 1, ethanol precipitated^82^, resuspended in 25 μL water, then electroporated into 100 μL aliquots of *B. burgdorferi* competent cells. The electroporated cells were immediately recovered in 6 mL BSK-II medium, and incubated overnight, after which selection in liquid culture and in semisolid BSK-agarose plates was performed. When needed, non-clonal, liquid-selected populations of transformants were plated in semisolid BSK-agarose for clone isolation. For each B31-derived clone, we confirmed the plasmid complement by multiplex PCR^83^. Plasmid complement was not determined for clones derived from strains other than B31 due to the absence of a complete, standardized PCR primer set for those strains. To confirm correct homologous recombination of suicide vectors, total genomic DNA was isolated from the *B. burgdorferi* clones using the Qiagen DNeasy Blood & Tissue Kit, and diagnostic PCR was performed to confirm insertion or deletions within the targeted locus, as well as correct recombination of the flanking sequences. The genetic manipulations of the chromosome at the *par* and *smc* loci are depicted schematically in Extended Data Fig. 8. Insertion of *parS^P1^* cassettes at the *phoU*, *uvrC*, and *lptD* loci occurred in intergenic regions between two convergently transcribed genes. The same strategy was used to insert the *parS^P1^* cassette in *B. burgdorferi* plasmids. For the cp32 plasmids, the *parS^P1^* cassette was inserted into the transposon sequence in transposon mutant backgrounds for which the transposon was mapped to have inserted in an intergenic region flanked by convergently transcribed genes^28^. As a result, the likelihood of coding sequence or promoter disruption by the genetic changes was minimized. An exception is lp25, where the cassette was inserted in the scar generated upon deletion of the nonessential gene *bbe02*.

### Mouse-tick transmission studies

#### Ethics statement

All animal work was performed according to the guidelines of the National Institutes of Health, *Public Health Service Policy on Humane Care and Use of Laboratory Animals* and the United States Institute of Laboratory Animal Resources, National Research Council, *Guide for the Care and Use of Laboratory Animals*. Protocols were approved by the Rocky Mountain Laboratories, NIAID, NIH Animal Care and Use Committee. The Rocky Mountain Laboratories are accredited by the International Association for Assessment and Accreditation of Laboratory Animal Care (AAALAC). All efforts to minimize animal suffering were made.

#### Experimental mouse-tick infection studies

Mouse infections were conducted with female RML mice, an outbred strain of Swiss-Webster mice reared at the Rocky Mountain Laboratories breeding facility. Five mice per strain were inoculated intraperitoneally (4 x 10^4^ spirochetes) and subcutaneously (1 x 10^4^ spirochetes), with the number of injected spirochetes pre-determined by Petroff-Hausser counting. Mouse infection was confirmed 3 weeks post-injection by isolation of *B. burgdorferi* from ear biopsies in BSK-II medium containing appropriate antibiotics.

Larval *Ixodes scapularis* were purchased from Oklahoma State University. *I. scapularis* were maintained between feeds at ambient light and temperature in bell jars over potassium sulfate-saturated water. Approximately 100 naïve *I. scapularis* larvae were fed to repletion per infected mouse. Acquisition and retention of *B. burgdorferi* by larval ticks was assessed 1 week after drop-off, and spirochete load was determined through mechanical disruption and plating. Two naïve mice per strain were fed upon by 15-20 infected *I. scapularis* nymphs. The number of *B. burgdorferi* in nymphs was assessed prior to feeding and 10 days after drop-off through mechanical disruption and plating. Mouse infection was confirmed 5 weeks post-nymphal tick feeding by isolation of *B. burgdorferi* from ear biopsies in BSK-II medium containing appropriate antibiotics.

### Tick midgut cryopreservation

Tick midguts were dissected and fixed on ice using 4% formaldehyde solution in PBS, washed thrice for 5 min with cold PBS, infiltrated with 15% sucrose in PBS at 4°C, then with 30% sucrose in PBS at 4°C, then with a 1:1 mix of 30% sucrose in PBS and Optimal Cutting Temperature (OCT) compound (Tissue-Tek), and finally with pure OCT compound. The tissue was then frozen in OCT compound using liquid-nitrogen-cooled isopentane, then stored at −80°C and shipped on dry ice. Thin, 10-μm sections were cut using a cryostat.

### Microscopy

*B. burgdorferi* culture density was measured by dilution of a culture in a C-Chip disposable hemocytometer (INCYTO) and direct counting of the cells under darkfield illumination obtained using a Nikon Eclipse E600 microscope equipped with a 40X 0.55 numerical aperture (NA) Ph2 phase-contrast air objective and darkfield condenser optics.

For fluorescence microscopy imaging, BSK-II-grown *B. burgdorferi* strains were spotted onto a 2% agarose-PBS pad^26,84^, covered with a no. 1.5 coverslip, then imaged using Nikon Eclipse Ti microscopes equipped with 100X Plan Apo 1.45 NA phase-contrast oil objectives, Hamamatsu Orca-Flash4.0 V2 CMOS cameras, and either a Sola LE light source or a Spectra X Light engine (Lumencor). The microscopes were controlled by Nikon Elements software. The following Chroma filter cubes were used to acquire the fluorescence images: DAPI, excitation ET395/25x, dichroic T425lpxr, emission ET460/50m; GFP: excitation ET470/40x, dichroic T495lpxr, emission ET525/50m; mCherry/TexasRed, excitation ET560/40x, dichroic T585lpxr, emission ET630/75m. DNA staining was obtained by incubating the culture for 15 min with Hoechst 33342 (Molecular Probes) at a final concentration of 1 μg/mL.

Tick midgut sections (see above) were processed as follows. The slides supporting the sections were brought up to RT, washed thrice with PBS, permeabilized for 30 min at RT using 0.2 % Triton X-100 in PBS, blocked for 30 min at RT using 5% BSA (w/v) in PBS, 0.1 % Tween-20, stained with RFP-Booster ATTO594 (ChromoTek) diluted 1:200 in blocking buffer (see above), washed thrice with PBS for 5 min each, and stained with Hoechst 33342 1:1000 in PBS for 10 min. The stained sections were mounted in PBS under a No 1.5 coverslip and were imaged on an OMX BLAZE microscope system (GE). Images in the DAPI, GFP, and mCherry channels were acquired at 100 nm z intervals. The image stacks were then deconvolved and registered using the Softworks software.

### Image analysis

Cell outlines were generated using phase-contrast images and the Oufti analysis software package^85^, with the following parameters: Edgemode, LOG; Dilate, 2; openNum, 3; InvertImage, 0; ThreshFactorM, 0.985; ThreshMinLevel, 0; EdgeSigmaL, 1; LogThresh, 0. Cell outlines were curated as follows: (i) outlines of cell debris were manually removed; (ii) outlines of cells that curled on themselves, crossed other cells, or were partially outside the field of view were also manually removed; (iii) partial outlines of cells were manually extended when feasible; (iv) outlines were manually added in some cases where automated outline generation failed; and (v) outlines were manually split for cells displaying clear phase profile dips around midcell, which indicated that the cytoplasmic cylinders of the daughter cells had separated yet remained linked by an outer membrane bridge^26^. This was confirmed by visual inspection of the fluorescence signal(s), as phase profile midcell dips often corresponded to a dip in the fluorescence signal. For this reason, throughout the manuscript, the term cell refers to an individual cytoplasmic cylinder. Lastly, the “Refine All” function of the Oufti software was ran, a final curation removed improper outlines, and the fluorescence signal was added to the cell outline file in Oufti. For signal quantification and intensity profile generation, we used the Oufti background fluorescence subtraction and intensity profile generation features. Image visualization was done using FiJi software^86^, GraphPad Prim software, and Adobe Illustrator.

To detect fluorescent spots, the Modified_Find_Irregular_Spots.m MATLAB function^87^ was used, with the following parameters: fitRadius: 5, edgeDist: 2.5, centerDist: 1; peakRadius: 3; shellThickness: 1; quantileThreshold: 0.3. The value of the intensityRatioThreshold parameter was determined individually for each dataset and fluorescence channel. For each intensityRatioThreshold value empirically tested, visual inspection of the accuracy of spot detection was performed on a subset of the cells using the VisualizeSpotDetection.m function. Once an appropriate intensityRatioThreshold value was identified, all the cell outlines were visually inspected and the ones that displayed under- or over-detection of spots were manually removed, yielding a curated cell list. Then, spots were identified and added to this final cell list using the add_spots_to_cellList.m routine, while the data was exported into a table format using export_to_table.m and extract_field.m routines. A summary of the imaging datasets is provided in Table S2, Worksheet 1. Each dataset was given a unique dataset identifier. Thus, replicate datasets of a given strain will have different identifiers. This summary includes the following information for each dataset: strain number, replicate number, relevant treatments, culture density at the time of image acquisition, number of cells in the dataset, and intensityRatioThreshold values used to detect the fluorescent spots. The *oriC*:*terC* ratio was calculated from microscopy data as follows: (i) the *oriC* and *terC* copy number per cell was plotted as a function of cell length in GraphPad Prism; (ii) a linear fit of the data was performed, and the slope of each fit was extracted; and (iii) the slope of the *oriC* fit was divided by the slope of the *terC* fit.

To quantify the extent to which the distribution of the *oriC* spots along the length of the cell deviates from an equidistant distribution, we performed the following analysis steps for each cell. First, we measured all distances between adjacent *oriC* spots, or DM (for distance measured). A cell with n spots will generate n-1 DM values. Next, in silico and for each cell individually, we equidistantly redistributed the *oriC* spots within the length of that cell, using the calculate_distance_ratio.m routine. This analysis assumes that each *oriC* spot is separated from the adjacent *oriC* spot or from the adjacent cell end (for the first and last *oriC* spot in a given cell, respectively) by the same distance. In this scenario, the distance DE between adjacent, equidistantly redistributed spots for a given cell is:

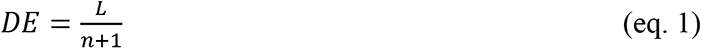

where L is the length of the cell and n is the number of *oriC* spots in the cell. Then, for each pair of adjacent *oriC* spots, we calculated a distance ratio, DR, defined as the ratio of the measured distance between two adjacent spots and the value this distance would have if all the spots in the cell were equidistantly spaced.

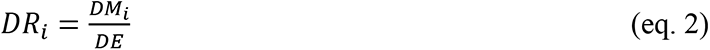

Therefore, for a perfectly equidistant distribution of *oriC* spots, all distance ratios are exactly 1. Finally, to assess how the *oriC* spots of a given cell deviate from equidistant spacing, we calculated a deviation from uniform spacing (or DUS) metric defined as the mean per cell of the absolute values of the deviation of the DR values from 1:

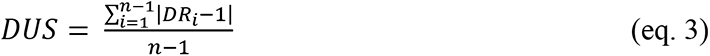

Please note that in a cell with n spots, there will be n-1 distances between adjacent spots. The data shown in Fig. 5I-K and 6E shows the distribution of DUS values within population of cells of the indicated strains.

We also simulated a random distribution of *oriC* copies inside the cells using the simulate_distance_ratio.m routine. For each analyzed cell (see above), this routine randomly redistributes each cell’s *oriC* copies along the length of that cell, then extracts the distances between adjacent randomly redistributed *oriC* spots and calculates the DR and DUS values as above.

### Sample growth for next-generation sequencing

A summary of the ChIP-seq samples analyzed and reported in this study is given in Table S2, Worksheet 2. For each strain and replicate, we provide information on culture growth conditions, inoculation and harvesting dates, and harvesting culture densities. The samples were prepared as follows. Exponentially growing *B. burgdorferi* strains were used to inoculate 250 mL BSK-II cultures. After two or three days of growth the cultures were fixed by addition of 95 mL 37% formaldehyde (Sigma-Aldrich #F8775) followed by incubation with rocking for 30 min at RT. Formaldehyde was then quenched by addition of 18 mL of 2.5 M glycine followed by incubation with rocking for 5 min at RT. The samples were chilled on ice for 10 min, transferred to conical centrifuge tubes and pelleted at 4°C using a 30 min, 4,300 x g spin in an Allegra X-14R centrifuge (Beckman Coulter) equipped with a swinging bucket SX4750 rotor. The pellet was resuspended in 30 mL ice-cold HN buffer (50 mM NaCl, 10 mM HEPES, pH 8.0)^88^, then pelleted at 4°C and 10,000 x g for 10 min in a fixed angle FX6100 rotor. The pellet was resuspended in 1.5 mL final cold HN buffer and pelleted once more at 4°C and 10,000 x g for 10 min. Finally, the pellet was resuspended in 500 μL ice-cold ChIP buffer A (12.5 mM Tris, 12.5 mM EDTA, 62.5 mM NaCl, 25% w/v sucrose, pH 8.0), frozen in a dry ice-ethanol bath, stored at −80°C.

### Chromatin immunoprecipitation-sequencing

The frozen *B. burgdorferi* cells were thawed on ice. One hundred microliters of cells were used to prepare DNA samples for whole genome sequencing (see below), and 400 μL were processed for chromatin immunoprecipitation-sequencing (ChIP-seq) as described previously^89^. Briefly, the fixed cells were lysed using lysozyme at 4 mg/mL final concentration. Crosslinked chromatin was sheared to an average size of 250 bp by sonication using a Qsonica Q800R2 water bath sonicator. The lysate was precleared using Protein A magnetic beads (Fisher 45-002-511) and was then incubated with anti-GFP antibodies^90^ or anti-mCherry antibodies^64^ overnight at 4°C. The next day, the lysate was incubated with Protein A magnetic beads for 1 h at 4°C. After washes and elution, the immunoprecipitate was incubated at 65°C overnight to reverse the crosslinks. The DNA was next treated with RNase A, proteinase K, extracted with phenol:chloroform:isoamyl alcohol (25:24:1), resuspended in 100 µL of buffer EB (Qiagen), and used for library preparation with the NEBNext UltraII kit (E7645). The library was sequenced using Illumina NextSeq500 at Indiana University’s Center for Genomics and Bioinformatics.

### Whole genome sequencing

For whole genome sequencing (WGS), 100 μL aliquots of the ChIP samples were pelleted, resuspended in 100 µL of TE (50mM Tris pH 8.0, 10 mM EDTA) containing 1 µL of proteinase inhibitor (Sigma P8340) and 6 µL of Ready-Lyse lysozyme (Epicentre, R1810M), and incubated at 37°C for 1.5 h. SDS was added to the cell suspension to a final concentration of 1% to solubilize the chromatin. A volume of 150 μL of TES (50mM Tris pH 8.0, 10 mM EDTA, 1% SDS) was added to the solution and the cell lysate was incubated at 65°C to reverse the crosslinks. After reversal of crosslinks, the WGS samples were processed and sequenced in the same way as ChIP-seq samples above. WGS of B31-derived strain S9 (see Table S1) was performed on cells grown and treated similarly to the ones used for the ChIP experiments described above.

Strain CJW_Bb523 was grown in 40 mL BSK-II medium to a density of ∼3-6 x 10^7^ cells/mL. The culture was pelleted at 4,000 x g for 10 min in the swinging bucket rotor, resuspended in 1 mL cold HN buffer, and re-pelleted. Genomic DNA was then extracted using DNeasy Blood and Tissue Kits (Qiagen) following the manufacturer’s recommendation for Gram-negative bacteria. Library preparation and whole-genome sequencing were done by the Yale Center for Genome Analysis on a NovaSeq6000 instrument with 2 x 150 bp read length.

### Sequence mapping and analysis

The sequencing reads for ChIP-seq and WGS were mapped to the combined *B. burgdorferi* genome (NCBI GCA_000008685.2_ASM868v2) using CLC Genomics Workbench (CLC Bio, Qiagen). Sequencing reads from each ChIP and WGS samples were normalized by the total number of reads. The WGS results were used as the “input” control for ChIP-seq samples. For marker frequency analysis, the reads corresponding to the *oriC* and *terC* regions were averaged over a 30 kbp span. For the plasmids, the reads were averaged over the entire size of each plasmid. The ChIP enrichment (ChIP/input) and the locus ratios were plotted and analyzed using R scripts, which are available from Dr. Xindan Wang upon request.

For the whole genome sequencing of strain CJW_Bb523, the sequencing reads were processed using Trimmomatic^91^ (parameter = PE, -phred33, -baseout, ILLUMINACLIP:2:30:10, TRAILING:20, MINLEN: 36) and around 8 million reads were mapped to *B. burgdorferi* B31 genome using Bowtie2^92^ (parameter = --non-deterministic). HTSeq^93^ was used to obtain the genomic coverage. The normalized count for each plasmid was calculated as the sum of reads at every base pair divided by the plasmid size. For *oriC*, we only considered reads mapped to nucleotide position 443,037-473,267 (∼30 kb region, similar to the average size of *Borrelia* plasmids) on the main chromosome.

### DNA fluorescence in situ hybridization (FISH)

*B. burgdorferi* cells were washed and resuspended in 1x PBS prior to the FISH experiments in order to remove contaminants from the BSK-H/BSK-II media. Cells were placed on poly-L-lysine coated wells that were outlined on coverslips with a hydrophobic pen (Super PAP pen, Invitrogen #008899). Prior to use, the coverslips were cleaned as described previously^39^. Once applied to the poly-L-lysine coated wells, *B. burgdorferi* cells were fixed in 4% formaldehyde in PBS for 5 min at RT. Cells were then washed 3 times in PBS. Cells were permeabilized using 400 μg/mL lysozyme in GTE buffer (50 mM glucose, 25 mM Tris, 10 mM EDTA, pH 8.0), then washed 3 times with PBS. A hybridization buffer was adapted from a previous report^94^. Briefly, 1 g dextran sulfate was dissolved in 5 mL dH2O, then 3530 μL formamide, 10 mg *E. coli* tRNA, 1 mL 20x SSC buffer (175.3 g/L NaCl, 88.2 g/L sodium citrate, pH 7.0), and 80 μL of 25 mg/mL BSA were added. The hybridization buffer was filtered and stored at −20°C. Cells, after fixation and permeabilization, were pre-hybridized in hybridization buffer containing 1 mg/mL RNase A for 30 min. Then, the cells were denatured on a heat block at 94°C for 5 min. First, the cells were placed on the heat block for 1 min in pre-hybridization solution, which was then replaced with hybridization solution, containing 200 nM probe. The Stellaris locked nucleic acid oligonucleotide FISH probe (sequence 5’-GAATAAGTAAAAGTGGTTTAG-3’, labeled with the dye 6FAM) was synthesized at Biosearch Technologies. This probe recognized a highly repetitive sequence present in 176 copies on plasmid lp21 of strain B31 and in the right subtelomeric region of the chromosome of strain 297^18,95^. The large number of repeats ensured that a strong fluorescent signal could be achieved despite using a single fluorescent probe. Strain B31e2^96^, which no longer harbors plasmid lp21 of the parental strain B31, served as a negative control as it lacks the target sequence of FISH probe. Hybridization proceeded at RT for 2 h. Wells were then washed as follows: 3 x 10 min washes with 40% (wt/vol) formamide in 2x SSC, 2 x 5 min washes in 2x SSC, and 3x washes in 1x PBS. Prior to imaging, 1 mg/mL DAPI in SlowFade® Gold Antifade Mountant (Thermo Fisher #S36937) was applied to each well.

### RNA isolation and qRT-PCR

Exponentially growing *B. burgdorferi* cultures were diluted to 2×10^5^ cells/mL in 30 mL BSK-II, grown in duplicate for 2 days, then RNA was extracted and quantified as previously described^75^. The cells were pelleted using a 10 min centrifugation at 4,300 x g in a swinging bucket rotor. The pellet was resuspended in 400 μL buffer HN^88^ (50 mM NaCl, 10 mM HEPES, pH 8.0). RNA was stabilized using the RNAprotect bacteria reagent (Qiagen), extracted using enzymatic lysis and proteinase K digestion (protocol 4 in the RNAprotect bacteria reagent manual), and purified using the RNeasy minikit (Qiagen). DNA was removed using the Turbo DNA-free kit (Thermo Fisher Scientific). RNA was quantified using the Kapa SYBR Fast one-step qRT-PCR mastermix kit (Roche) using 10 ng total RNA per reaction, in duplicate. One additional reaction was performed for each sample without the reverse transcriptase and confirmed that the measured amplification was not due to DNA contamination of the RNA samples. The primers used for amplification of the *parZ* transcript were 5’-CCCCCTATTTTAAAAACCGAAG-3’ and 5’-TAATGGTTTGCGCGTATCC-3’. The primers used for amplification of the control *recA* transcript were previously described^19^. The amount of the *parZ* transcript was normalized to the level of the *recA* transcript and then expressed for each sample relative to mean levels measured in strain CJW_Bb488, as previously described^97^.

### Phylogenetic analyses

Protein sequences were retrieved from the NCBI databases by Standard Protein BLAST searches using *B. burgdorferi* proteins as queries in the NCBI BLASTP web-based platform. For Borreliaceae ParZ sequence alignments and phylogenetic tree building and for *Brachyspira*, *Leptospira*, and *Treponema* ParB sequence alignments, only non-redundant RefSeq protein sequences were analyzed. For *Leptospira,* the analysis of ParB sequences was limited to those encoded by chromosome I. For ParZ-like phylogenetic tree building, all BLAST hits retrieved using *B. burgdorferi* ParZ as query were utilized. The BLAST search was performed across the three domains of life and among virus sequences. Sequences were aligned and the phylogenetic trees were built using the Geneious sequence analysis software.

### Statistics and data visualization

Statistical analyses were performed in GraphPad Prism software. Data visualization was achieved using MATLAB software, GraphPad Prism software, and Adobe Illustrator. Statistical summaries for all figure panels that involve data quantification are provided in Table S2, Worksheet 3, and include the following information: strain used in the figure, number of imaging datasets analyzed, and total number of cells for the combined replicates.

### Data and material availability

The ChIP-Seq and WGS data discussed in this study are deposited in the NCBI Gene Expression Omnibus platform and are available through GEO Series accession number GSE202255 (https://www.ncbi.nlm.nih.gov/geo/query/acc.cgi?acc=GSE202255). The accession numbers for each sample can be found in Table S2 Worksheet 2. The plasmids and strains generated in this study will be made available by the Jacobs-Wagner laboratory upon publication of the study. The image analysis code developed as part of this study was deposited in the publicly accessible Github code repository (https://github.com/JacobsWagnerLab/published).

### Plasmid construction methods

Plasmids are listed in Table S1, Worksheet 2. Oligonucleotide primers used in the plasmid generation process are listed in Table S1, Worksheet 3.

I. *Shuttle vectors*

**pBSV2G_P_0826_-mCherry^Bb^-ParB.** The following fragments were assembled (using intermediary constructs) between the SacI and PstI sites of plasmid pBSV2G_2: a) the promoter of *B. burgdorferi* gene *bb0826* (P*_0826_*)^79^, corresponding to bp 535523-535703 of strain B31’s chromosome, PCR-amplified with primers NT115 and NT116 and digested with SacI and BamHI; b) the *mcherry^Bb^* sequence^79^, amplified with primers NT100 and NT101 and digested with KpnI and BamHI; c) *parB* (*bb0434*), amplified from strain B31’s genomic DNA with primers NT230 and NT232 and digested with SalI and PstI.

**pBSV2G_P_0826_-RBS-ParZ-msfGFP^Bb^.** The following fragments were assembled (using intermediary constructs) between the SacI and KpnI and between the PstI and HindIII sites of plasmid pBSV2G_2, respectively: a) promoter P*_0826_*^79^, flanked by SacI and BamHI restriction enzyme sites; b) *parZ* (*bb0432*), PCR-amplified from strain B31’s genomic DNA with primers NT363 and NT364 and digested with BamHI and KpnI; c) the *msfgfp^Bb^* sequence^79^, PCR-amplified with primers NT159 and NT160 and digested with PstI and HindIII.

**pKFSS1_P_0826_-mCherry^Bb^-ParB.** P*^0826^*-*mcherry^Bb^*-*parB* was moved from pBSV2G_P_0826_-mCherry^Bb^-ParB into pKFSS1_2 using SacI and HindIII.

**pKFSS1_P_0826_-msfGFP^Bb^-ParB.** *msfgfp^Bb^* was PCR amplified using primers NT341 and NT162, digested with BamHI and KpnI and ligated into the BamHI/KpnI backbone of pKFSS1_P_0826_-mCherry^Bb^-ParB.

**pBSV2G_P_0031_-mCherry^Bb^-ParB^P1^**. The following fragments were assembled, through intermediates, between the SacI/KpnI, and XbaI/HindIII sites of pBSV2G_2: promoter P*_0031_*^79^, obtained from strain B31’s genomic DNA by amplification with primers NT111 and NT112 and digestion with SacI and BamHI; *mcherry^Bb^*, PCR amplified with NT100 and NT101 and digested with BamHI and KpnI; and *parB^P1^*, obtained by de novo gene synthesis at Genewiz flanked by XbaI and HindIII restriction enzyme sites. The *parB^P1^* gene encodes the P1 plasmid ParB protein minus its N-terminal peptide sequence (ParB^ΔN30^)^98^, which was codon-optimized for translation in *B. burgdorferi* as previously described^79^ and was deposited at GenBank under accession number ON321895.

**pBSV2G_P_0031_-msfGFP^Bb^-ParB^P1^.** *msfgfp^Bb^* was PCR amplified using primers NT161 and NT162, digested with BamHI and KpnI, and cloned into the BamHI/KpnI sites of pBSV2G_P_0031_-mCherry^Bb^-ParB^P1^.

**pBSV2B_P_0826_-mCherry^Bb^-ParB.** P*_0826_*-*mcherry^Bb^*-*parB* was moved from pBSV2G_P_0826_-mCherry^Bb^-ParB into pBSV2B using SacI and HindIII digestion.

**pBSV2B_r(P_0031_-msfGFP^Bb^-ParB^P1^)_P_0826_-mCherry^Bb^-ParB.** P*_0031_*-*msfgfp^Bb^*-*parB^P1^* was released from pBSV2G_P_0031_-msfGFP^Bb^-ParB^P1^ as a SacI/FspI fragment and was ligated into the SacI/BsrBI backbone of pBSV2B_P_0826_-mCherry^Bb^-ParB.

**pBSV2G_P_smcL_-mCherry^Bb^-Smc.** The following fragments were sequentially assembled within the multicloning site of plasmid pBSV2G_2: *mcherry^Bb^*, PCR-amplified using primers NT100 and NT101 and digested with BamHI and KpnI; a DNA fragment of 804 bp upstream of the ttg START codon of *smc* (*bb0045*), PCR*-* amplified with primers NT217 and NT219 and digested with SacI and BamHI; and a DNA fragment encoding *smc* (*bb0045*), PCR-amplified using primers NT221 and NT222 and digested with SalI and PstI.

II. *. Suicide vectors*

Note on nomenclature: “pKI” signifies a suicide vector generated to mediate knock-in of a gene of interest into the *B. burgdorferi* genome. While most of these plasmids are based on pCR2.1, some are based on other backbones. The antibiotic resistance used for selection is noted in the name of the construct (i.e. pKIKan, pKIGent, or pKIStrep) and is driven by default by the P*_flgB_* promoter. When, instead, the P*_flaB_* promoter was used to drive antibiotic resistance gene expression, this is noted in the name of the plasmid (e.g., pKIKan(P_flaB_)).

**pKIGent.** This plasmid, obtained through a series of intermediate constructs, contains the following features: a) a Δ*aphI*Δ*bla* backbone of plasmid pCR2.1^75^, flanked by HindIII and XbaI restriction enzyme sites; b) a P*_flgB_*-*aacC1*-*flaB*t cassette flanked by SpeI and SacII restriction enzyme sites. This cassette is flanked by two multicloning sites, namely HindIII-KpnI-SacI-BamHI-SpeI-XmaI and SacII-EcoRI-PstI-NotI-XhoI-SphI-XbaI. To generate this cassette, the following fragments were assembled through intermediates: (i) P*_flgB_*-*aacC1*^77^ was cloned in between the XmaI and the SacII sites of the backbone; (iii) *flaB*t, the flagellin transcriptional terminator, was generated by annealing primers NT350 and NT351 and ligating them into the SpeI/XmaI sites of the backbone. This inactivated the original XmaI site of the backbone but generated a new one downstream of *flaB*t.

**pKIGent_parS^P1^.**This plasmid differs from pKIGent, in that the *parS^P1^* sequence^99^ was synthesized de novo at Genewiz and cloned into the pUC57Amp backbone. It was then PCR-amplified using primers NT165 and NT166, digested with SpeI and XmaI, and inserted into the SpeI/XmaI sites of the backbone. The resulting clone had a mutation in the *parS^P1^* sequence that was corrected by site-directed mutagenesis using primers NT215 and NT216.

**pKIGent_parS^P1^_phoU.** *B. burgdorferi* B31 chromosomal region from nucleotide 38650 to 40720 was PCR-amplified with primers NT175 and NT176 and digested with SacI and SpeI. The chromosomal region from nucleotide 40721 to 42797 was PCR-amplified with primers NT177 and NT178 and digested with PstI and XhoI. The two fragments were inserted into the SacI/SpeI and PstI/XhoI sites of pKIGent_parS^P1^, respectively.

**pKIGent_parS^P1^_cp26.** *B. burgdorferi* B31 cp26 region from nucleotide 20585 to 22602 was PCR-amplified with primers NT203 and NT204 and digested with SacI and BamHI. The cp26 region from nucleotide 22603 to 24632 was PCR-amplified with primers NT205 and NT206 and digested with PstI and XhoI. The two fragments were inserted into the SacI/BamHI and PstI/XhoI sites of pKIGent_parS^P1^, respectively.

**pKIGent_parS^P1^_uvrC.** *B. burgdorfe*ri strain B31’s chromosomal region from nucleotide 474180 to 476218 was PCR-amplified with primers NT267 and NT268 and digested with SacI and SpeI. The region from nucleotide 476251 to 478279 was PCR-amplified with primers NT269 and NT270 and digested with PstI and XhoI. The two fragments were inserted into the SacI/SpeI and PstI/XhoI sites of pKIGent_parSP1, respectively.

**pΔparBS.** *B. burgdorferi* chromosomal region between nucleotides 448842 and 450913 was PCR-amplified with primers NJ99 and NJ100 and digested with BamHI and XmaI. The region from nucleotide 452017 to 453998 was PCR-amplified using primers NJ97 and NJ98 and digested with PstI and XhoI. The two fragments were inserted into the BamHI/XmaI and PstI/XhoI sites of pKIGent, respectively.

**pΔparBS(Kan).** The P*_flaB_-aphI* cassette from pKIKan(P_flaB_) was excised using PstI and XmaI and inserted into the PstI/XmaI sites of pΔparBS.

**pKIGent_par.** The *B. burgdorferi* chromosomal region from nucleotide 448842 to 450913 was PCR-amplified with primers NJ99 and NJ100 and digested with BamHI and XmaI. The region between nucleotides 451133 and 453037 was PCR-amplified with primers NJ101 and NJ102 and digested with PstI and XhoI. The two fragments were inserted into the BamHI/XmaI and PstI/XhoI sites of pKIGent, respectively.

**pKIGent_parS^P1^_lp17.** The 1.3 kbp BamHI/FspI fragment of pKIGent_parS^P1^ was ligated with the 2.9 kbp BamHI/NaeI backbone of pKK81.

**pΔparAZ.** Nucleotides 447274 through 449320 of strain B31 chromosome were PCR-amplified using primers NT530 and NT531, digested with BamHI and XmaI, and ligated into the BamHI/XmaI backbone of pKIGent_par.

**pΔparZ.** Nucleotides 448134 through 450172 of the B31 chromosome were PCR-amplified using primers NT528 and NT529, cut with BamHI and XmaI, and ligated into the BamHI/XmaI backbone of pKIGent_par.

**pΔpar.** Nucleotides 447274 through 449320 of the B31 chromosome were PCR-amplified using primers NT530 and NT531, digested with BamHI and XmaI, and ligated into the BamHI/XmaI backbone of pΔparBS.

**pKIGent_parS^P1^_lp28-3.**Nucleotides 1853 to 3891 of plasmid lp28-3 were PCR-amplified using primers NT487 and NT488 and digested with KpnI and BamHI. Nucleotides 3890 to 5944 of plasmid lp28-3 were PCR-amplified using primers NT489 and NT490 and digested with SacII and XhoI. These fragments were cloned into the corresponding sites of pKIGent_parS^P1^.

**pKIGent_parS^P1^_lp28-2.**Plasmid lp28-2 was PCR-amplified using primers NT483 and NT484 and digested with KpnI and SpeI. Plasmid lp28-2 was PCR-amplified using primers NT485 and NT486 and digested with SacII and XhoI. These fragments were cloned into the corresponding sites of pKIGent_parS^P1^.

**pKIGent_parS^P1^_lp25.** The P*_flgB_*-*aacC1*-*parS^P1^* cassette was PCR-amplified from pKIGent_parS^P1^ using primers NT509 and NT524, cut with PacI and MluI, and cloned into the PacI/MluI backbone of pKbeKan.

**pKIGent_parS^P1^_lp38.** Nucleotides 20973 through 23014 of plasmid lp38 were PCR-amplified using primers NT499 and NT500, then digested with KpnI and SpeI. Nucleotides 23014 through 25091 of plasmid lp38 were PCR-amplified using primers NT501 and NT502, then digested with SacII and XhoI. The fragments were ligated into the corresponding sites of pKIGent_parS^P1^.

**pKIGent_parS^P1^_lp36**. Plasmid lp36 was PCR-amplified with primers NT495 and NT496, digested with KpnI and BamHI, then cloned into the KpnI/BamHI sites of pKIGent_parS^P1^. The resulting plasmid was PCR-amplified using primers MRS_17 and MRS_18 and Gibson-assembled with a PCR product obtained by amplification of plasmid lp36 using primers MRS_19 and MRS_20.

**pKIKan.** This plasmid was obtained through a series of intermediate constructs, and in a manner similar to pKIGent, except that is has an *aphI* kanamycin resistance gene instead of an *aacC1* gentamicin resistance gene under the control of P*<u>_flgB_</u>*. It contains the following features: a) the Δ*aphI*Δ*bla* backbone of plasmid pCR2.1^75^, flanked by HindIII and XbaI restriction enzyme sites; b) a P*_flgB_*-*aphI*-*flaB*t cassette flanked by SpeI and SacII restriction enzyme sites that are part of two multicloning sites: HindIII-KpnI-SacI-BamHI-SpeI-XmaI and SacII-PstI-NotI-XhoI-SphI-XbaI. To generate this cassette, the following fragments were assembled through intermediates: (i) P*_flgB_*, flanked by SacII and NdeI sites; (ii) *aphI*, PCR-amplified from pBSV2 using primers NT534 and NT535, digested with NdeI and XmaI, and cloned together with P*_flgB_* between the SacII and XmaI sites of the backbone; and (iii) *flaB*t, generated by annealing primers NT350 and NT351 and ligating them into the SpeI/XmaI sites of the backbone. This inactivated the original XmaI site of the backbone but generated a new one downstream of *flaB*t.

**pKIKan(P_flaB_).** P*_flaB_* was PCR-amplified from pBSV2G_P_flaB_-mCerulean^Bb^ using primers NT577 and NT578, digested with SacII and NdeI, and ligated into the SacII/NdeI backbone of pKIKan.

**pKIKan_parS^P1^.** P*_flgB_*-*aphI-flaB*t was moved from pKIKan into the backbone of pKIGent_parS^P1^ as an XmaI/PstI fragment.

**pKIKan(P_flaB_)_parS^P1^.** The P*_flaB_*-*aphI*-*flaB*t cassette was moved from pKIKan(P_flaB_) into the backbone of pKIGent_parS^P1^ as an XmaI/PstI fragment.

**pKIKan_parS^P1^_uvrC.** P*_flgB_*-*aphI*-*parS^P1^* was excised from pKIKan_parS^P1^ using SpeI and PstI and inserted into the SpeI/PstI backbone of pKIGent_parS^P1^_uvrC, where it replaced P*_flgB_*-*aacC1*-*parS^P1^*.

**pKIGent_parS^P1^_lp28-4.**The pKIGent_parS^P1^ backbone was PCR-amplified using primers MRS_46 and MRS_31. Plasmid lp28-4 was PCR-amplified using primers MRS_49 and MRS_53, and MRS_51 and MRS_52, respectively. The P*_flgB_*-*aacC1*-*parS^P1^* cassette was PCR-amplified from pKIGent_parS^P1^ using primers MRS_32 and MRS_33. The four PCR products were Gibson assembled together.

**pKIGent_parS^P1^_lp54.** pABA01 was PCR-amplified using primers MRS_24 and MRS_29. The P*_flgB_*-*aacC1*-*parS^P1^* cassette was PCR-amplified from pKIGent_parS^P1^ using primers MRS_25 and MRS_26. The two PCR products were Gibson assembled.

**pΔparS.** A region between nucleotides 449961 and 451998 of strain B31’s chromosome was PCR-amplified using primers NT625 and NT626, digested with BamHI and XmaI, and inserted into the BamHI/XmaI sites of plasmid pΔparBS.

**pKIKan(P_flaB_)_par.** The P*_flaB_-aphI-flaB*t cassette from pKIKan(P_flaB_) was excised using PstI and XmaI and inserted into the PstI/XmaI sites of pKIGent_par.

**pKIKan(P_flaB_)_ParZ-msfGFP^Bb^.** *msfgfp^Bb^* was PCR-amplified with primers NT643 and NT644. Plasmid pKIKan(P_flaB_)_par was PCR-amplified with primers NT645 and NT646. The two fragments were Gibson-assembled.

**pKIKan(P_flaB_)_mCherry^Bb^-ParB.** *mcherry^Bb^* was PCR-amplified with primers NT629 and NT630. Plasmid pKIKan(P_flaB_)_par was PCR-amplified with primers NT631 and NT632. The two fragments were Gibson-assembled.

**pKIGent_parS^P1^_lp28-1.** P*_flgB_-aacC1_parS^P1^* was PCR-amplified from pKIGent_parS^P1^ using primers NT762 and NT763. The backbone of p28-1::flgBp-aacC1 was PCR-amplified using primers NT764 and NT765. The two fragments were Gibson-assembled together.

**pKIKan(P_flaB_)_ParA-msfGFP^Bb^.** *msfgfp^Bb^* was PCR-amplified using primers NT633 and NT634. *parA* was PCR-amplified using primers NT636 and NT768. *parZ* was PCR-amplified from pKIKan(P_flaB_)_par using primers NT766 and NT635. The suicide vector backbone was PCR-amplified from pKIKan(P_flaB_)_par using primers NT769 and NT767. The four PCR products were Gibson-assembled together.

**pKIKan(P_flaB_)_ParZ-msfGFP^Bb^_ΔparA.** Site-directed mutagenesis was performed on plasmid pKIKan(P_flaB_)_ParZ-msfGFP^Bb^ using primers NT623 and NT624.

**pKIKan(P_flaB_)_ParA-msfGFP^Bb^_ΔparZ.** Site-directed mutagenesis was performed on plasmid pKIKan(P_flaB_)_ParA-msfGFP^Bb^ using primers NT778 and NT779.

**pKIKan(P_flaB_)_ΔparBS_ParA-msfGFP^Bb^_ΔparZ.** The BamHI/XmaI insert of plasmid pKIKan(P_flaB_)_ParA-msfGFP^Bb^_ΔparZ was ligated into the BamHI/XmaI backbone of plasmid pΔparBS(Kan).

**pKIKan(P_flaB_)_ΔparBS_ParA-msfGFP^Bb^.** The BamHI/XmaI insert of plasmid pKIKan(P_flaB_)_ParA-msfGFP^Bb^ was ligated into the BamHI/XmaI backbone of plasmid pΔparBS(Kan).

**pKIKan(P_flaB_)_ΔparBS_ParZ-msfGFP^Bb^.** The BamHI/XmaI insert of plasmid pKIKan(P_flaB_)_ParZ-msfGFP^Bb^ was ligated into the BamHI/XmaI backbone of plasmid pΔparBS(Kan).

**pKIKan(P_flaB_)_mCherry^Bb^-ParB_ΔparAZ.** The BamHI/XmaI insert of plasmid pΔparAZ was ligated into the BamHI/XmaI backbone of plasmid pKIKan(P_flaB_)_mCherry^Bb^-ParB.

**pKIKan(P_flaB_)_ParA-msfGFP^Bb^_ParZ^ΔN20^.** Site-directed mutagenesis was performed on pKIKan(P_flaB_)_ParA-msfGFP^Bb^ using primers NT1020 and NT1021.

**pKIGent_parS^P1^_lp21_V2.**The backbone of pKIGent_parS^P1^ was PCR-amplified with primers NT800 and NT801. The P*_flgB_-aacC1_parS^P1^* cassette was PCR-amplified from pKIGent_parS^P1^ with primers NT796 and NT797. Nucleotides 253 through 1114 of the B31 plasmid lp21 were PCR-amplified using primers NT794 and NT795. Nucleotides 1115 through 2628 of the B31 plasmid lp21 were PCR-amplified using primers NT798 and NT799. The four PCR products were Gibson-assembled.

**pKIGent_parS^P1^_lptD**. The backbone of pKIGent_parS^P1^ was PCR-amplified with primers NT824 and NT825. The P*_flgB_-aacC1_parS^P1^* cassette was PCR-amplified from pKIGent_parS^P1^ with primers NT820 and NT821. Nucleotides 895077 through 896572 of the B31 chromosome were PCR-amplified using primers NT818 and NT819. Nucleotides 896573 through 898070 of the B31 chromosome were PCR-amplified using primers NT822 and NT823. The four PCR products were Gibson-assembled.

**pKIGent_mCherry^Bb^-Smc.**The following fragments were Gibson-assembled: 1) nucleotides 43811 through 45353 of the B31 chromosome fused downstream of and in frame to *mcherry^Bb^*, controlled by the *smc* native promoter, obtained by PCR-amplification of plasmid pBSV2G_P_smcL_-mCherry^Bb^-Smc with primers NT1006 and NT1007; 2) the gentamicin cassette of pKIGent_parS^P1^_phoU, obtained by PCR-amplification with primers NT1008 and NT1009; 3) nucleotides 45376 through 46530 of the B31 chromosome, obtained by PCR-amplification with primers NT1010 and NT1011; and 4) the suicide vector backbone of plasmid pΔparA(kan), obtained by PCR-amplification with primers NT1012 and NT1013.

**pΔsmc(gent).** The following fragments were Gibson-assembled: 1) nucleotides 42016 through 43530 of the B31 chromosome, obtained by PCR amplification with primers NT960 and NT961; 2) the gentamicin cassette of pKIGent_parS^P1^_phoU, obtained by PCR amplification with primers NT962 and NT963; 3) nucleotides 45332 through 46857 of the B31 chromosome, obtained by PCR amplification with primers NT964 and NT965; and 4) the suicide vector backbone of plasmid pΔparA(kan), obtained by PCR amplification with primers NT966 and NT967.

**pKIGent_ΔparBS_ParZ-msfGFP^Bb^_ΔparA.** The gentamicin resistance cassette was excised from pΔparB as a SacII/XmaI fragment and inserted into the SacII/XmaI backbone of plasmid pKIKan(P_flaB_)_ΔparBS_ParZ-msfGFP^Bb^_ΔparA.

**pKIKan(P_flaB_)_ΔparBS_ParZ-msfGFP^Bb^_ΔparA.** The BamHI/XmaI insert of plasmid pKIKan(P_flaB_)_ParZ-msfGFP^Bb^_ΔparA was ligated into the BamHI/XmaI backbone of plasmid pΔparBS(Kan).

**pKIStrep(P_flaB_)_parS^P1^.** *aadA* was PCR-amplified from pJSB252 using with primers NT756 and NT757. The backbone of the pKIKan(P_flaB_)_parS^P1^ plasmid was PCR-amplified using primers NT758 and NT759. The two fragments were Gibson-assembled with each other.

**pKIStrep_parS^P1^_Tn.**pGKT was digested with EagI, and the resulting 1.8 kbp fragment was purified and ligated, yielding plasmid pTG, which was then PCR-amplified using primers NT690 and NT691. P*_flaB_*-*aadA*_*parS^P1^* was PCR-amplified from pKIStrep(P_flaB_)_parS^P1^ using primers NT692 and NT693. The two PCR products were Gibson assembled with each other. The resulting plasmid was PCR amplified with NT784 and NT785 and the resulting PCR product was digested with AvrII and self-ligated.

## EXTENDED DATA FIGURE LEGENDS

**Extended Data Figure 1.**
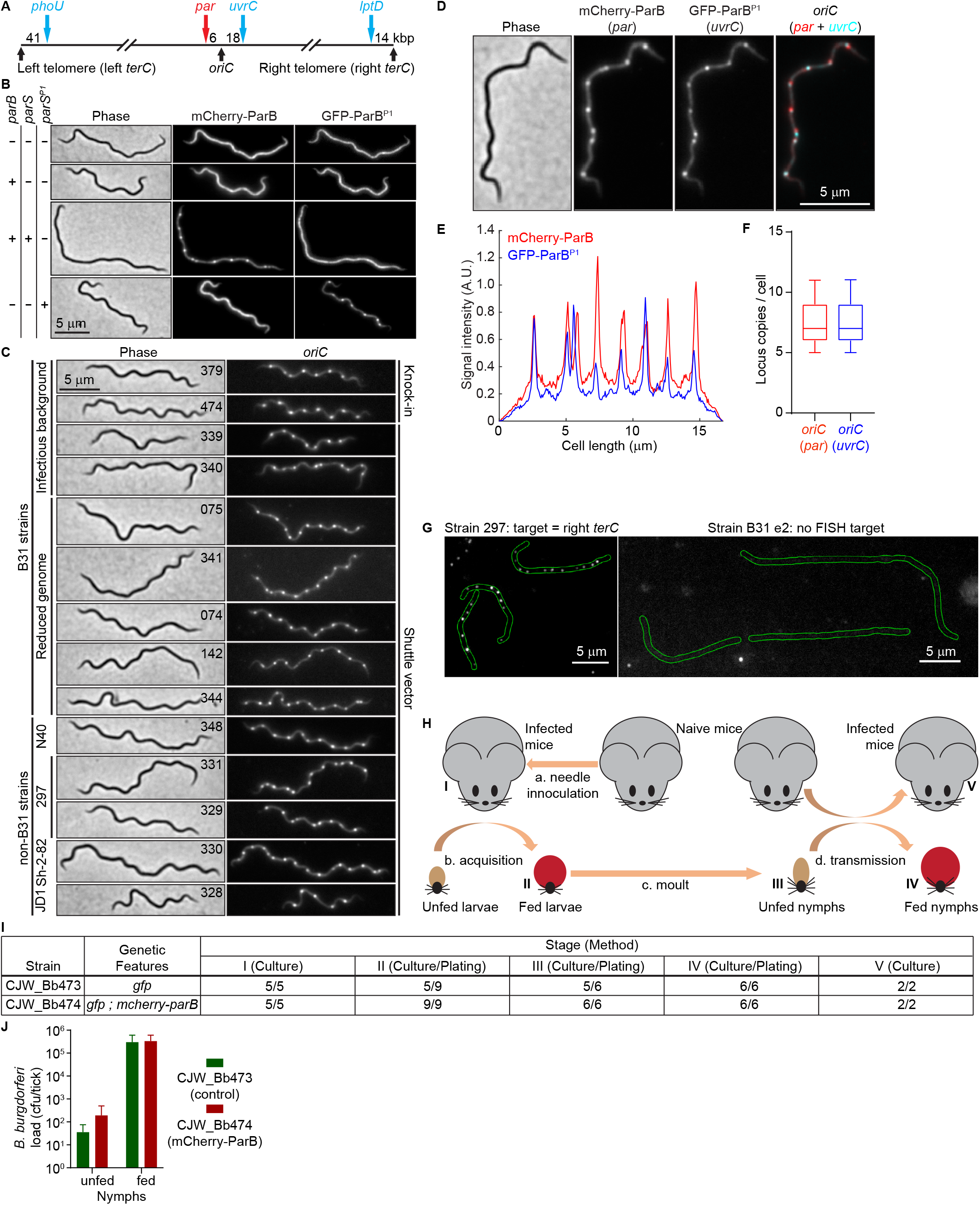
*B. burgdorferi* cells carry multiple chromosome copies. **A.** Schematic of chromosomal loci localized in this study (not drawn to scale). *oriC* was localized either by labeling the *par* locus through the expression of fluorescently tagged ParB (red) or by insertion of *parS^P1^* downstream of *uvrC*, which is located 52% along the length of the chromosome, followed by expression of GFP-ParB^P1^. The telomeres were labeled by insertion of *parS^P1^* at the *phoU* or *lptD* loci, which are located at 5% or 98% along the length of the chromosome, respectively, followed by expression of GFP-ParB^P1^. Distances between the labeled DNA loci and the *oriC* or *terC* loci, are shown in kilobase pairs (kbp). **B.** Images showing that mCherry-ParB and GFP-ParB^P1^ specifically recognize their cognate *parS* sites. mCherry-ParB and GFP-ParB^P1^ were expressed from the same shuttle vector (see methods). The strains are, from top to bottom: CJW_Bb211, CJW_Bb534, CJW_Bb532, and CJW_Bb533. Presence of endogenous *parB* or *parS*, and chromosomal insertion of *parS^P1^* are indicated at the left. **C.** Localization of mCherry-ParB or GFP-ParB at *oriC* regions in various strain backgrounds. Tagged ParB was expressed either by knock-in of *mcherry-parB* at the gene locus or in trans, from a shuttle vector. Strain CJW_Bb142 expressed *msfgfp-parB* from a shuttle vector. Strain backgrounds are shown at the left. The CJW_Bb number of each strain is listed on the phase-contrast image. **D.** Images of a cell of strain CJW_Bb205 showing the *oriC* region co-labeled by expression of mCherry-ParB, which binds to the endogenous *parS* site located within the *par* locus, and GFP-ParB^P1^, which binds to the *parS^P1^* sequence introduced at the *oriC*-proximal *uvrC* locus, as shown in (A). **E.** Fluorescence intensity profiles along the cell length for the cell shown in (C). **F.** Boxplot showing the number of *oriC* copies per cell based on the labeling of the *par* locus (red) or of the *uvrC* locus (blue) in strain CJW_Bb205. Shown are the mean of the data (middle line), the 25 to 75 percentiles of the data (box), and the 2.5 to 97.5 percentiles of the data (whiskers). **G.** Images showing DNA fluorescence in situ hybridization (FISH) staining of the repetitive sequence found within the right telomere of the chromosome of strain 297. Strain B31e2, which does not contain this repetitive sequence, serves as a negative control. Cell outlines are in green. **H.** Schematic of the experimental workflow used to test the transmission of *B. burgdorferi* strains CJW_Bb473 and CJW_Bb474 between ticks and mice. Roman numerals depict the stages at which infection of mice or colonization of ticks by *B. burgdorferi* was assessed. **I.** Summary of infection or colonization readouts as assayed at the stages depicted in (H). Assay methods are given for each stage. Shown are numbers of positive animals (mice or ticks) over numbers of assayed animals. **J.** Plot showing *B. burgdorferi* loads in nymphs prior to nymphal stage feeding (unfed) or 10 days after nymphal feeding drop-off (fed).

**Extended Data Figure 2.**
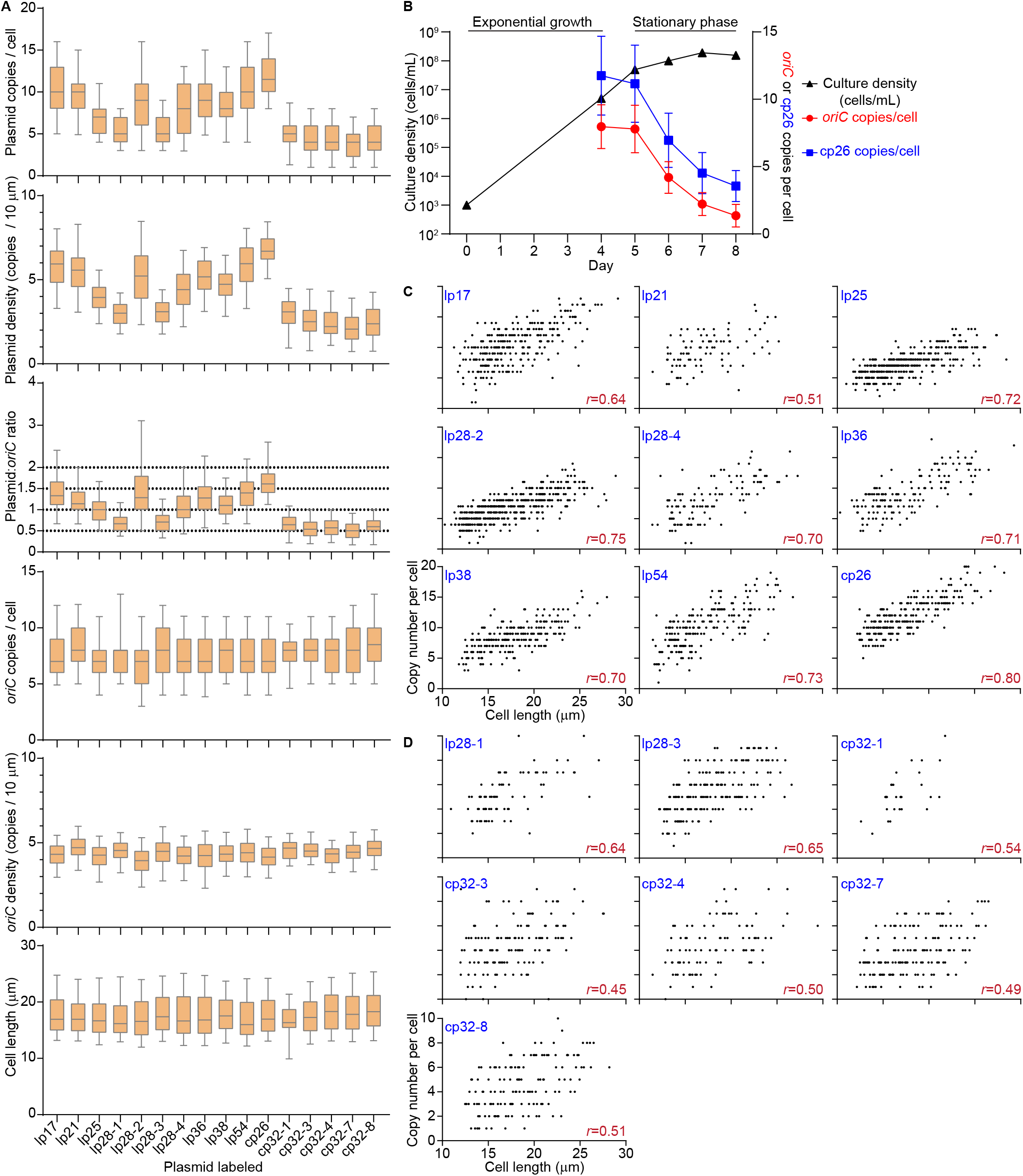
*B. burgdorferi* contains multiple copies of its endogenous plasmids. **A.** Boxplots showing the quantification of various characteristics (plasmid copies per cell; plasmid copies per 10 μm of cell length; plasmid to *oriC* ratios; *oriC* copies per cell; *oriC* copies per 10 μm of cell length, and cell length) for strains in which an endogenous plasmid is labeled by insertion of *parS^P1^* and expression of GFP-ParB^P1^, while *oriC* is labeled by expression of mCherry-ParB. Strains are, from left to right: CJW_Bb207, CJW_Bb526, CJW_Bb274, CJW_Bb489, CJW_Bb271, CJW_Bb241, CJW_Bb325, CJW_Bb272, CJW_Bb261, CJW_Bb326, CJW_Bb203, CJW_Bb501, CJW_Bb515, CJW_Bb517, CJW_Bb516, and CJW_Bb518. Selected images for each of these strains are provided in Fig. 2A. Shown are the mean of the data (middle line), the 25 to 75 percentiles of the data (box), and the 2.5 to 97.5 percentiles of the data (whiskers). **B.** An exponentially growing culture of strain CJW_Bb203, in which *oriC* is labeled by expression of mCherry-ParB and cp26 is labeled using the GFP-ParB^P1^/parS^P1^ was diluted to 10^3^ cells/mL, then imaged daily from day 4 through day 8 of growth in culture. Shown is the *oriC* copy number per cell (red, mean ± standard deviation), the cp26 copy number per cell (blue, mean ± standard deviation) and the culture density (black, in cells/mL) at the indicated times. **C.** Correlations between plasmid copy number per cell and cell length in a subset of the strains described in (A) and Fig. 2A. The analyzed plasmid is listed in blue, while the Spearman’s correlation coefficient *r* is in burgundy. **D.** Correlations between plasmid copy number per cell and cell length in the remaining strains described in (A) and Fig. 2A and not included in (B). These plasmids have fewer copies per cell. The analyzed plasmid is listed in blue, while the Spearman’s correlation coefficient *r* is in burgundy.

**Extended Data Figure 3.**
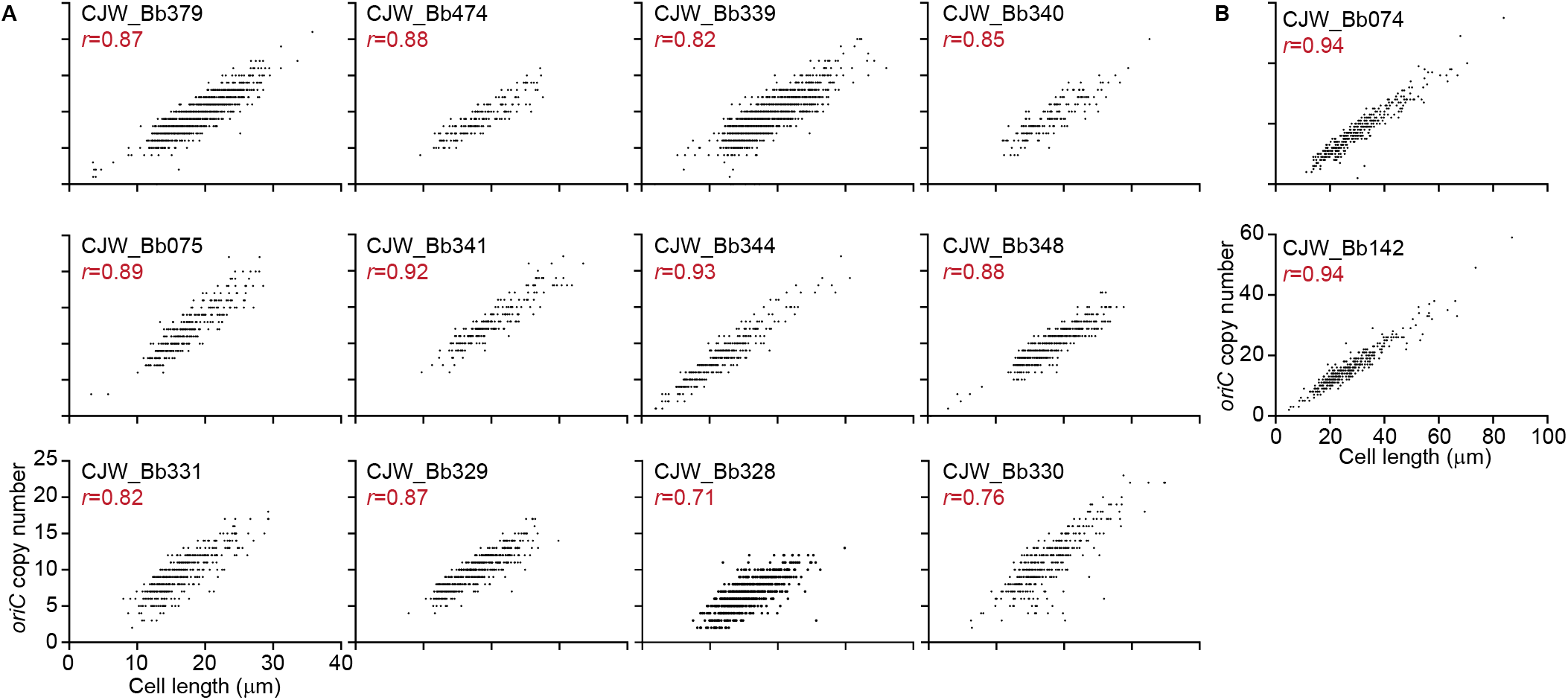
Chromosome copy number correlates with cell length. **A.** Correlations between *oriC* copy number per cell and cell length in the indicated strains, which are also shown and analyzed in Fig. 1B and Extended Data Fig. 1C. *r*, Spearman’s correlation coefficient. **B.** Same as in (A), except for strains CJW_Bb074 and CJW_Bb142. These strains have longer characteristic cell lengths, which is reflected in the range used for the x-axis.

**Extended Data Figure 4.**
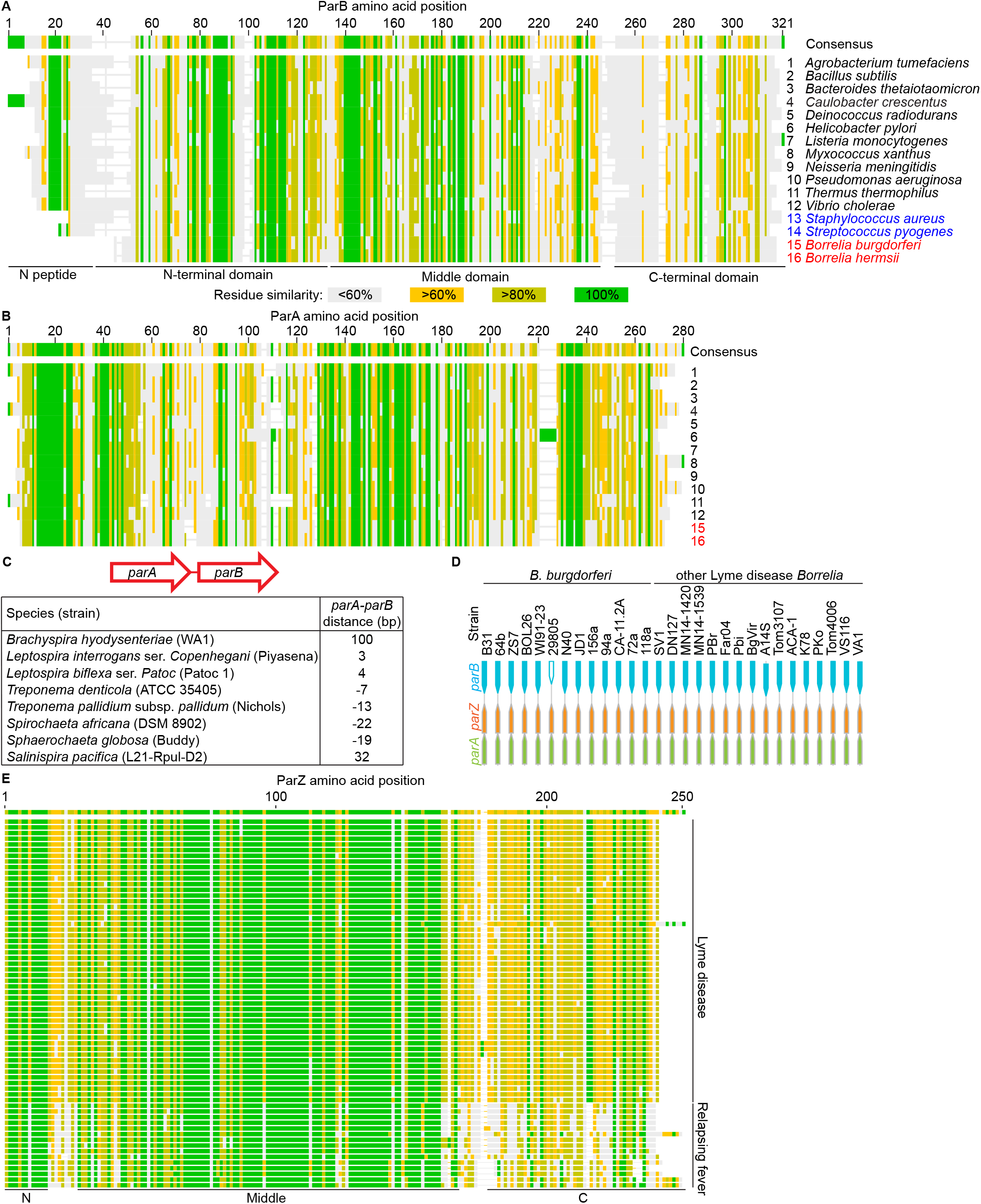
Phylogenetic analyses of Par proteins. **A.** Alignment of the indicated chromosomally expressed ParB sequences. ParB domains are highlighted at the bottom. **B.** Alignment of indicated chromosomally expressed ParA sequences. The numbers at the right indicate the same species as those listed in (A), at the right. **C.** Table showing the distance in base pairs (bp) between *parA* and *parB* homologs in the *par* loci of representative spirochete bacteria. *parA* and *parB* are found in the same orientation with a short genomic distance separating the two genes, which is suggestive of an operon structure. A negative value indicates overlap of the coding regions of the two genes. **D.** Organization of the *par* loci of the indicated Lyme disease spirochete strains as visualized using the BorreliaBase genome browser^100^. **E.** Alignment of 65 Borreliaceae ParZ sequences. Putative ParZ domains are highlighted at the bottom. Sequences belonging to Lyme disease and relapsing fever spirochetes are marked at the right.

**Extended Data Figure 5.**
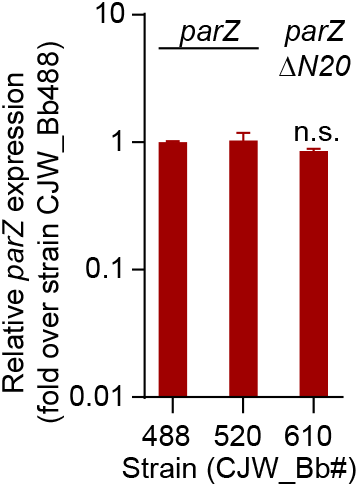
ParZ transcript levels. Plot showing relative mRNA levels for *parZ* and *parZ*Δ*N20* determined by qRT-PCR in the indicated strains.

**Extended Data Figure 6.**
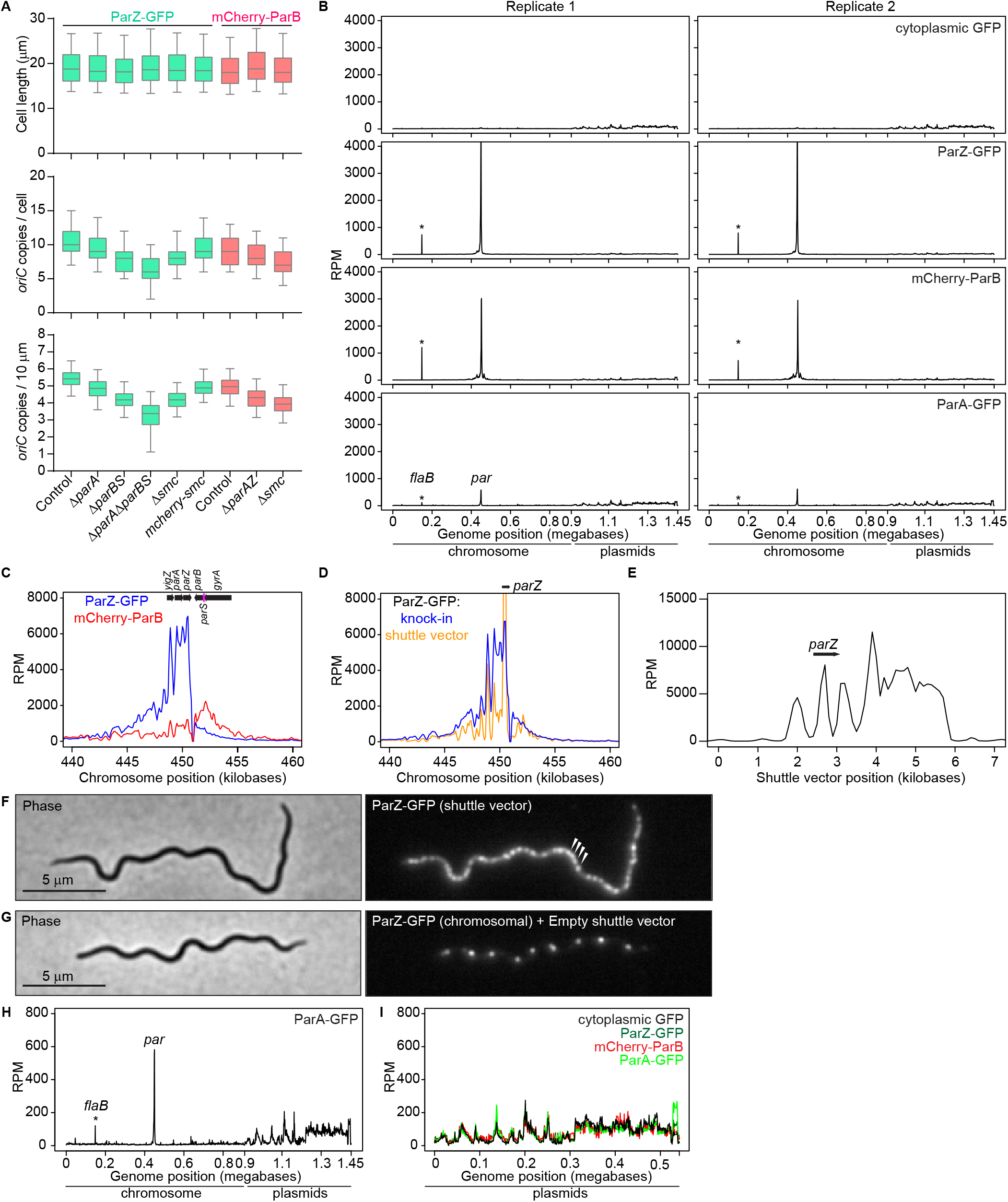
ParZ is a novel centromere-binding protein that controls oriC segregation. **A.** Boxplots comparing cell length, *oriC* copy numbers per cell, and *oriC* copies per 10 μm of cell length for control and mutant strains analyzed in Fig. 5 and 6. Shown are the mean of the data (middle line), the 25 to 75 percentiles of the data (box), and the 2.5 to 97.5 percentiles of the data (whiskers). The nature of the *oriC* label is listed at the top. Mutations introduced into the strains are listed at the bottom. From the left, the following strains were used: CJW_Bb378, CJW_Bb490, CJW_Bb524, CJW_Bb616, CJW_Bb603, CJW_Bb602, CJW_Bb379, CJW_Bb525, and CJW_Bb604. **B.** Whole genome ChIP-seq profiles for strains expressing free GFP (CJW_Bb473), ParZ-GFP (CJW_Bb378), mCherry-ParB (CJW_Bb379), or ParA-GFP (CJW_Bb488). The x-axis shows the chromosome coordinates followed by the concatenated endogenous plasmids of strain S9 in the order: lp28_3, lp25, lp28_2, lp38, lp36, lp28_4, lp54, cp26, lp17, lp28_1, cp32_1, cp32_3, cp32_4, cp32_6, cp32_7, cp32_8, cp32_9, and lp21. Two replicates are shown for each strain. No ChIP-seq peaks are seen in the free GFP control. Two peaks are seen in each of the other traces. The main peak is located at the *par* locus. Please see Fig. 5E-H for a detailed view of this *par* region of the traces, including in additional strains. The second peak is at the *flaB* locus (marked by a star in the figure) but does not represent specific binding of ParZ-GFP, mCherry-ParB or ParA-GFP to the *flaB* locus. Rather, it represents binding of these proteins to the P*_flaB_*-*aphI*-*flaB*t kanamycin resistance cassette that we inserted into the *par* locus during strain generation, in between *parZ* and *parB.* These bound P*_flaB_* and *flaB*t reads were then mapped to sequences within native *flaB* locus, to which they are identical. RPM, reads per million. **C.** ChIP-seq profiles of ParZ-GFP and mCherry-ParB binding to the *par* locus in strain CJW_Bb403, which expresses both protein fusions. RPM, reads per million. The dip in the trace between *parZ* and *parB* is due to read mapping to the reference B31 chromosomal sequence which does not contain the kanamycin resistance cassette present in the strains analyzed by ChIP-seq (see supplemental text). Same is true for (D). **D.** ChIP-seq profiles of ParZ-GFP binding to the *par* locus in strains CJW_Bb378, which has a *parZ-msfgfp* knock-in at the *parZ* gene locus, and CJW_Bb101, which carries *parZ-msfgfp* on a shuttle vector. Binding of ParZ-GFP to the *parZ* sequence (highlighted) likely appears higher in strain CJW_Bb101 because the strain contains more copies of the *parZ* sequence, located on the chromosome and on the shuttle vector, than in strain CJW_Bb378, where all *parZ* sequences are chromosomal. **E.** ChIP-seq profile of ParZ-GFP binding to the shuttle vector encoding *parZ-msfgfp* in strain CJW_Bb101. The *parZ* gene location is highlighted. Binding may be skewed to the right due to transcription as *parZ-msfgfp* and the adjacent antibiotic resistance cassette are oriented to the right. **F.** Images of a cell of strain CJW_Bb101. Arrowheads pinpoint four of the many densely packed ParZ-GFP puncta that can be detected in cells of this strain. **G.** Images of a cell of a strain CJW_Bb571 which expresses ParZ-GFP from the *parZ* gene locus and carries an empty shuttle vector. **H.** Magnified view of the ParA-GFP replicate 1 ChIP-seq trace shown in (B). **I.** ChIP-seq profiles showing an overlay of the landscape binding pattern of free GFP, ParZ-GFP, mCherry-ParB, and ParA-GFP to the concatenated endogenous plasmid sequences. Traces are those of Replicate 2 shown in (B).

**Extended Data Figure 7.**
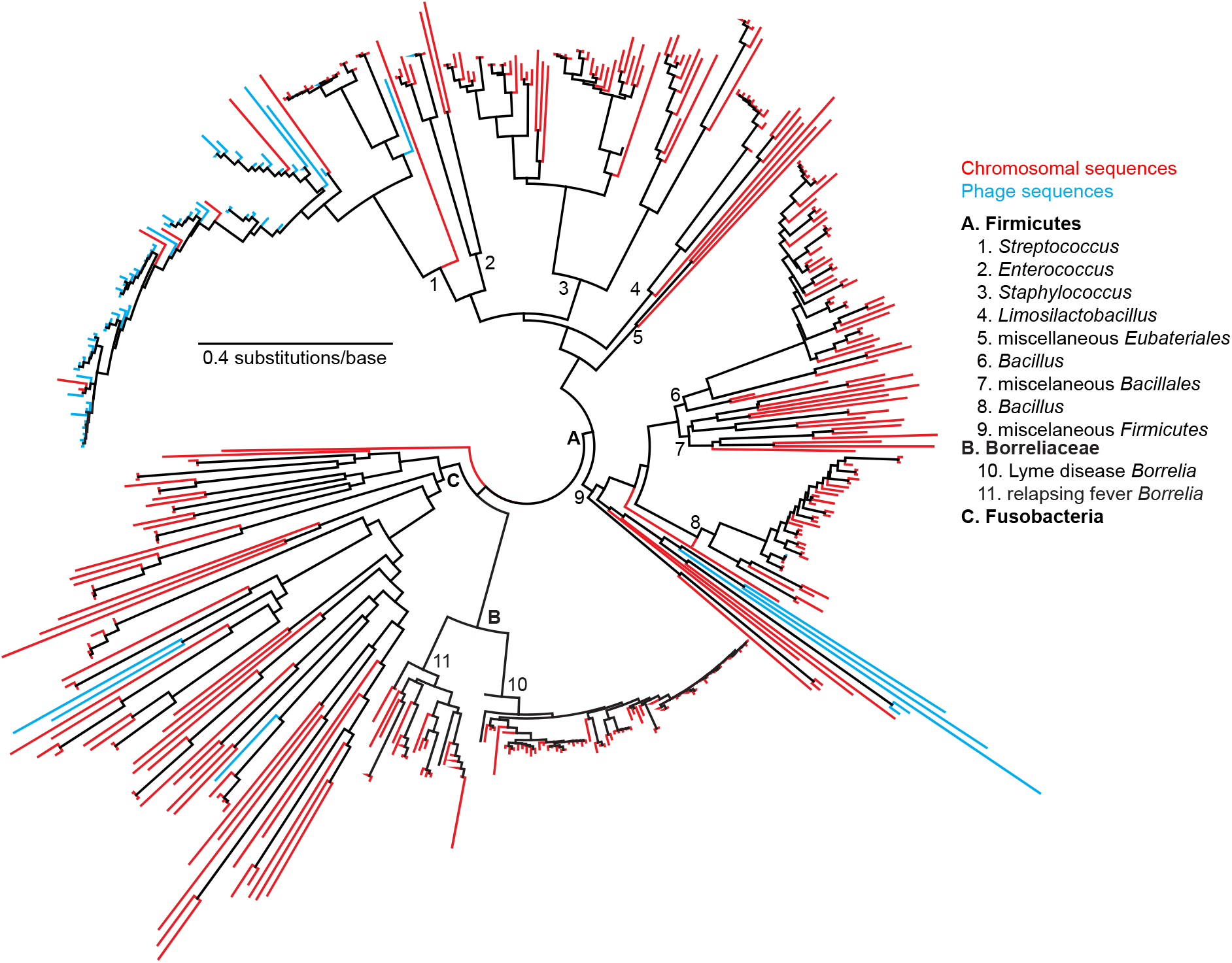
ParZ-like sequences can be found in Firmicutes, Fusobacteria, and their phages. Blast searches were performed using *B. burgdorferi* ParZ as bait. No hits were obtained among archaeal and eukaryotic proteins. Hits obtained among bacterial chromosome-encoded proteins are in red, while those obtained among bacteriophage-encoded proteins are in cyan. Letters highlight the Firmicutes and Fusobacteria phyla, or the Borreliaceae family, while the numbers highlight the indicated genera.

**Extended Data Figure 8.**
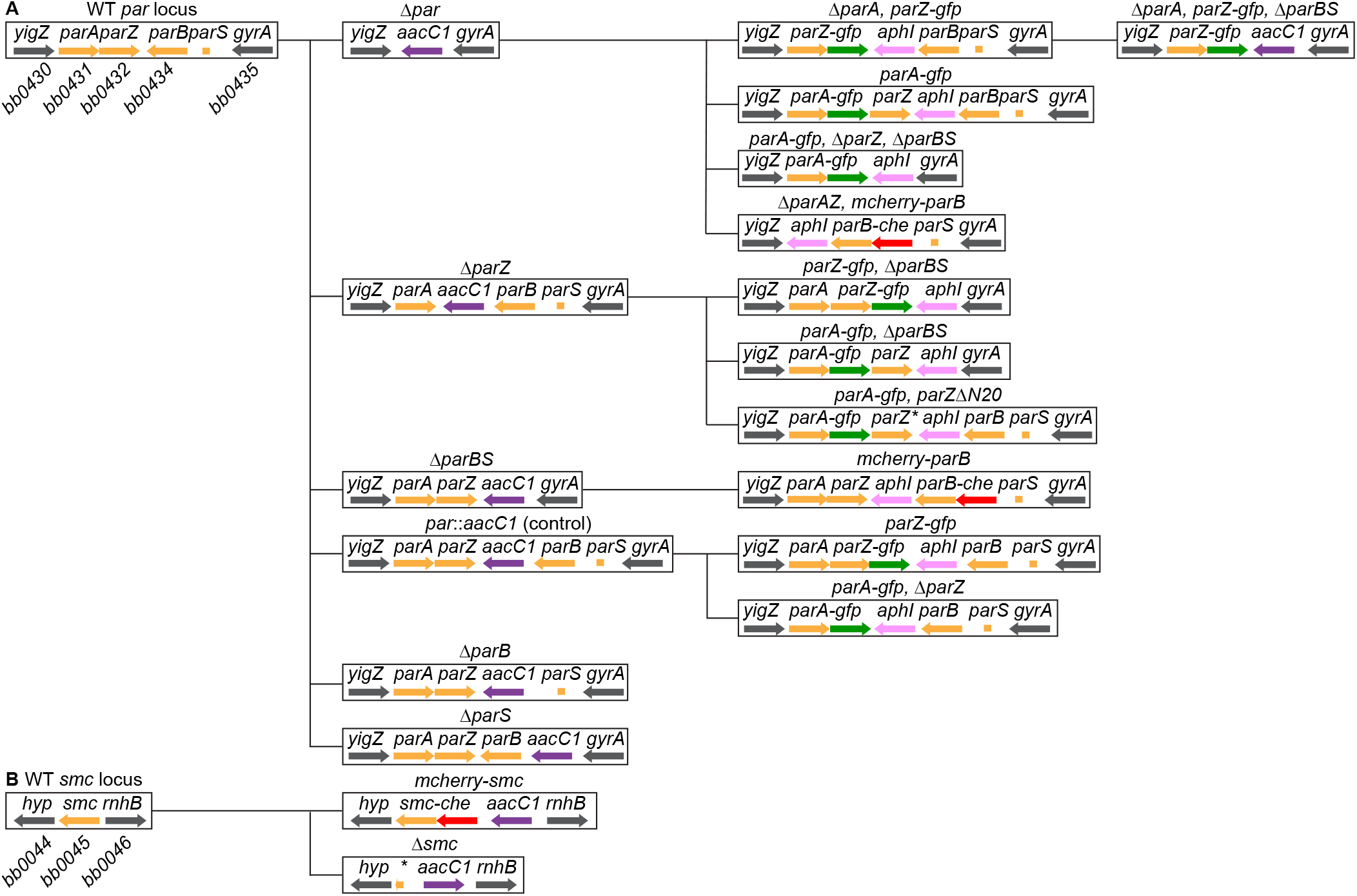
Schematic pedigree of genetic changes engineered at the *par* and *smc* loci. **A.** Depiction of genetic changes at the *par* locus. Genes affected by genetic modifications are in orange. Genes flanking the modified region and not affected by the changes are in gray. WT, wild type. *aacC1*, gentamicin resistance cassette. *aphI*, kanamycin resistance cassette. The promoters and transcriptional terminators present in the antibiotic resistance cassettes are not shown. Features are not drawn to scale. The lines starting from the WT locus at the left depict the order in which successive genetic modifications were introduced. *, *parZ*Δ*N20*. **B.** Same as in (A) but for the *smc* locus. *hyp*, gene *bb0044* of hypothetical function. *, a short sequence encoding the C-terminus of SMC was not deleted to avoid inactivating the promoter upstream of gene *bb0044*. A. and B. Please note that individual strains (see Table S1) may carry genetic modification at a single locus or multiple loci, including the chromosomal *phoU*, *uvrC*, or *lptD* loci, or plasmid-specific loci.

