## Supplemental text for "Polyploidy, regular patterning of genome copies, and unusual control of DNA partitioning in the Lyme disease spirochete"

**Table of contents……………………………………………………………………………..Page**

Specific labeling of *B. burgdorferi* DNA loci using endogenous and heterologous ParB/*parS* systems……………………………………………………………………………………….2

Detection of *oriC* loci in multiple *B. burgdorferi* strains………………………………………....3

Recapitulation of the mouse-tick transmission cycle using strain CJW_Bb474………………….4

Chromatin immunoprecipitation-sequencing (ChIP-seq) of Par proteins………………………...5

Evidence of ParZ interaction with its own coding region…………………………………………7

References……………………………………………………………………………………7

*Specific labeling of* B. burgdorferi *DNA loci using endogenous and heterologous ParB/*parS *systems*

ParB proteins specifically recognize their cognate *parS* sequence and spread onto adjacent DNA sequences[1-3]. Due to this property, expression of a fluorescent protein-tagged ParB protein leads to accumulation of its fluorescence into a diffraction-limited signal that pinpoints the subcellular location of the DNA locus that contains the *parS* sequence[4].

We have adapted this method for use in *B. burgdorferi*, whose chromosome contains a single predicted *parS* sequence[3] located within the *par* locus, 6 kilobases to the left of *oriC* (Extended Data Fig. 1A). We labeled this *parS* sequence either by replacing the native *parB* gene (*bb0434*) with the *mcherry-parB* translational fusion, yielding knock-in strains (Fig. 1A, Extended Data Fig. 1C, Table S1), or by driving expression of *mcherry-parB* or *msfgfp-parB* from a multi-copy shuttle vector (SV) using the weak promoter $P_{0826}$[5] (Extended Data Fig. 1C, Table S1). mCherry-ParB fluorescent foci formed only when *parS* was present on the *B. burgdorferi* chromosome (Extended Data Fig. 1B).

To label an additional *B. burgdorferi* locus, we first inserted the *parS* sequence of *E. coli* plasmid P1[2], hereafter referred to as $parS^{P1}$, into the *B. burgdorferi* genome. We then expressed an *msfgfp* fusion to the *parB* gene of plasmid P1 ($msfgfp\text{-}parB^{P1}$) from the same multi-copy shuttle vector that contained the *mcherry-parB* expression cassette (Table S1, Extended Data Fig. 1B). We drove expression of $msfgfp\text{-}parB^{P1}$ using the intermediate strength promoter $P_{0031}$[5]. The expressed GFP-ParB$^{P1}$ formed fluorescent puncta only when $parS^{P1}$ was also present in a given *B. burgdorferi* strain (Extended Data Fig. 1B), regardless of whether the chromosomal *parS* site was present or not (Extended Data Fig. 1B,D), confirming that labeling of the two *parS* sequences by their tagged cognate ParB proteins was independent and specific.

This conclusion was further strengthened by quantitative analyses of images of strain CJW_Bb205 (Extended Data Fig. 1D-F). In this strain, mCherry-ParB foci, which pinpoint the subcellular location of *par* loci, and GFP-ParB$^{P1}$ foci, which pinpoint the subcellular location of *uvrC* loci (Extended Data Fig. 1A), colocalized almost perfectly (Extended Data Fig. 1D-E). The *par* and *uvrC* loci are 24 kbp away from each other and 6 and 18 kbp away from *oriC*, respectively (Extended Data Fig. 1A). Thus, both labels approximate the subcellular location of *oriC*. Importantly, copy numbers of the *par* and *uvrC* loci were similar (Extended Data Fig. 1F).

*Detection of* oriC *loci in multiple* B. burgdorferi *strains*

We localized the *oriC* locus in several *B. burgdorferi* strains that were derived from the B31 isolate, which is the type strain, as well as from other isolates, namely N40, 297, Sh-2-82, and JD1. For the B31-derived strains, we used the B31-A3-68-Δ*bbe02* genetic background (strains S9 and K2, see Table S1), which is easily transformable and fully capable of completing the tick-mouse transmission cycle[6,7]. The S9 derivatives CJW_Bb379 and CJW_Bb474 both carry a replacement of the *parB* gene with an *mcherry-parB* fusion driven by the native *parB* promoter, and are therefore labeled as knock-in (KI) strains (Fig. 1B, Extended Data Fig.1C). CJW_Bb474 additionally expresses free GFP, driven by the $P_{flaB}$ promoter, and inserted into endogenous plasmid cp26 (Fig. 1A, Table S1). Strains CJW_Bb339 and CJW_Bb340 are also derived from the infectious K2 and S9 strains, respectively, but express *mcherry-parB* as a second *parB* copy, in trans, from a shuttle vector (Table S1). Strains CJW_Bb339, CJW_Bb340, CJW_Bb379, and

CJW_Bb474 each has an almost complete complement of endogenous plasmids. They only lack plasmids cp9, lp5, and lp56 (Table S1), which are also absent from the parental strains S9 and K2 and are not required for completion of the tick-mouse transmission cycle[6,8-10]. We therefore refer to these strains as having an infectious background, which we experimentally demonstrated for strain CJW_Bb474 (see below).

We determined that the other B31-derived strains have lost multiple endogenous plasmids (Table S1) during their generation and/or the generation of their parental strains[11-13]. At most, strain CJW_Bb075 carries 11 endogenous plasmids, while strain CJW_Bb344 only carries two endogenous plasmids, cp26 and cp32-3 (Table S1). They all expressed tagged ParB proteins (mCherry-ParB or GFP-ParB) from a shuttle vector (Table S1). Lastly, we localized *oriC* loci in several other *B. burgdorferi* strains, including the widely studied N40, 297, and JD1 isolate backgrounds (Fig. 1B, Extended Data Fig. 1C, Table S1). We did not determine the endogenous plasmid content of the clones derived from the non-B31 isolates as there are no available characterized sets of primers for multiplex PCR detection of the native plasmids of these strains.

*Recapitulation of the mouse-tick transmission cycle using strain CJW_Bb474*

Strain CJW_Bb474 was used to image the chromosomal copy number in the tick (Fig. 1H). Since this strain carries genetic modifications, it was important to assess whether it can reproduce the mouse-tick transmission cycle. Two modifications, inactivation of gene *bbe02* and constitutive expression of GFP from cp26, did not affect *B. burgdorferi*'s ability to complete its transmission cycle when previously tested in several strain backgrounds[6,7,9,10,14,15]. The third modification, replacement of *parB* with *mcherry-parB*, has not been previously tested. Expanded

Data Fig. 1H depicts our experimental setup. Mice were infected with *B. burgdorferi* by needle inoculation (step a). Naïve tick larvae were allowed to feed on these infected mice and thus to acquire *B. burgdorferi* (step b). These colonized larvae molted into unfed nymphs (step c), which were then allowed to feed on and transmit *B. burgdorferi* to naïve mice (step d). Infection of mice was confirmed by tissue biopsy culture in BSK-II medium (stages I and V). *B. burgdorferi* acquisition by, and stable colonization of, ticks were assessed in fed larvae, unfed nymphs, and fed nymphs (stages II through IV) by crushing ticks in BSK-II then using the resulting tick extracts to inoculate liquid BSK-II cultures or embedding them in semisolid BSK-agarose plates. Spirochete outgrowth in the BSK-II medium or colony formation in the BSK-agarose plates were deemed evidence that the ticks were colonized by *B. burgdorferi*. All the mice exposed to strain CJW_Bb474, as well as those exposed to the CJW_Bb473 control strain, which only expresses GFP from cp26, were successfully infected (Expanded Data Fig. 1I). All the ticks exposed to CJW_Bb474 were also infected, as were most of the ticks exposed to the CJW_Bb473 control (Expanded Data Fig. 1I). Additionally, spirochete loads in unfed nymphs were close to $10^2$ cfu/tick for both strains (Expanded Data Fig. 1J). These loads increased to above $10^5$ cfu/tick in fed nymphs assayed 10 days after completion of nymphal feeding (Expanded Data Fig. 1J). The spirochete burdens that we measured in unfed and fed nymphs are similar to those previously measured in ticks colonized with the parental strain S9[16-18]. These results indicate that strain CJW_Bb474 is fully capable of completing the mouse-tick transmission cycle.

*<u>Chromatin immunoprecipitation-sequencing (ChIP-seq) of Par proteins</u>*

We mapped the reads from our ChIP-seq experiments to a theoretical reference genome obtained by concatenating the sequences of the chromosomes and plasmids of strain B31 (Expanded Data

Fig. 6B). We did not include the cp9, lp5, or lp56 plasmid sequences as these plasmids are absent from all the strains we used in the ChIP-seq experiments. The strains also contain genetic modifications such as: the presence of an antibiotic resistance cassette introduced during strain generation, the presence of fluorescent protein-coding sequences, or the absence of deleted sequences in the mutant strains. To preserve our ability to compare mapped reads between strains, unless indicated otherwise, we mapped the reads to the reference sequence and visualized the results accordingly. As a result, mapping to the *par* locus reveals a dip in the trace between *parZ* and *parB* (Fig. 5E-H, 6C, and Expanded Data Fig. 6C-D). This reflects replacement of the intergenic region between *parZ* and *parB* with a $P_{flaB}$-*aphI-flaBt* kanamycin resistance cassette. The $P_{flaB}$ (promoter of the flagellin gene *flaB*) and *flaBt* (transcriptional terminator of the flagellin gene *flaB*) sequences that we inserted into the *par* locus as part of the kanamycin cassette are identical to those found at the native *flaB* locus of the B31 chromosome. As a result, mCherry-ParB, ParZ-GFP, and ParA-GFP binding to the *par* locus also involves binding to the $P_{flaB}$ and *flaBt* sequences within. While these reads map to the native *flaB* locus (Extended Data Fig. 6B,H), they do not reflect actual binding to that locus.

Additionally, we observed low-level, non-uniform mapping of ChIP-seq reads to *B. burgdorferi*'s endogenous plasmids (Expanded Data Fig. 6B,H). Since free GFP, ParZ-GFP, ParA-GFP, and mCherry-ParB pulldowns generated almost identical traces in this region (Expanded Data Fig. 6I), we believe these traces represent non-specific landscape binding.

*Evidence of ParZ interaction with its own coding region*

As ParZ-GFP formed puncta in the absence of *parA* or *parBS* (Fig. 5A), we hypothesized that the centromere-like sequence recognized by ParZ resides within the *parZ* gene itself. To test this hypothesis, we expressed ParZ-GFP from a shuttle vector that has about five-fold more copies than the chromosome[19-21]. In the resulting strain CJW_Bb101, ParZ-GFP bound the chromosomal *par* locus (Extended Data Fig. 6D), as expected, but also the shuttle vector itself (Extended Data Fig. 6E). The skewed spreading of ParZ-GFP on the shuttle vector to the right of *parZ* (Extended Data Fig. 6E) may be facilitated by the direction of transcription of *parZ-msfgfp* and of the antibiotic resistance gene *aacC1*, both of which are transcribed to the right. Consistent with the ChIP-seq results, cells of strain CJW_Bb101 had numerous ParZ-GFP puncta (Extended Data Fig. 6F). We presume these puncta represent both chromosomal *oriC* loci and individual copies of the shuttle vector. Consistent with the high copy number of the shuttle vector, the ParZ-GFP puncta in cells of strain CJW_Bb101 were far more numerous than those generated by chromosomally expressed ParZ-GFP (Fig. 5A and Extended Data Fig. 6G). Additionally, the presence of an empty shuttle vector in a strain expressing ParZ-GFP from the chromosome did not cause an increase in the density of the ParZ-GFP puncta (Extended Data Fig. 6G), ruling out recruitment of ParZ-GFP by the shuttle vector backbone.
